# ELITE: E3 Ligase Inference for Tissue specific Elimination: A LLM Based E3 Ligase Prediction System for Precise Targeted Protein Degradation

**DOI:** 10.1101/2025.11.05.686884

**Authors:** Sabyasachi Patjoshi, Sumit Madan, Holger Fröhlich

## Abstract

Targeted protein degradation (TPD) has transformed modern drug discovery by harnessing the ubiquitin-proteasome system to eliminate disease-driving proteins previously considered undruggable. However, current approaches predominantly rely on a narrow set of ubiquitously expressed E3 ligases, such as Cereblon (CRBN) and Von Hippel-Lindau (VHL), limiting tissue specificity, increasing the risk of systemic toxicity, and facilitating resistance. Here, we present an AI-driven framework for the rational identification of tissue-specific E3 ligases for precision-targeted degradation. The framework leverages protein language models trained on large-scale protein sequence corpora to generate contextual embeddings that capture structural and functional features relevant to E3-substrate compatibility. These representations are integrated with tissue-resolved protein-protein interaction data to predict biologically plausible, context-dependent ligase-target interactions and prioritize E3 ligases for disease-relevant tissues. The framework was evaluated across 26 individual tissue contexts and further assessed in a pooled all-tissue setting in which tissue identity was incorporated explicitly through a learned tissue embedding. Robustness across protein representations was examined using ProtBERT, ESM-2 650M, and ESM-C, with ESM-C providing the strongest overall predictive performance. The strong performance of ESM-C further suggests that it may provide an effective foundation for broader protein-protein interaction applications beyond E3-substrate prediction, including generic PPI prediction and host-pathogen interaction modeling. The approach was additionally benchmarked against recent sequence-based deep-learning methods for E3-substrate interaction prediction. Together, these evaluations demonstrate that tissue context can be incorporated effectively into protein foundation-model-based interaction prediction while maintaining strong performance across heterogeneous biological settings. By curating and harmonizing heterogeneous biological data specifically for tissue-aware E3-ligase identification, the proposed framework adapts general-purpose protein language models to a previously unexplored application in targeted protein degradation. This provides a scalable strategy for expanding the therapeutically accessible E3-ligase repertoire and advancing TPD toward greater tissue selectivity and precision medicine.

## Significance

Current computational strategies for targeted protein degradation (TPD) largely ignore biological context, relying on global E3–substrate datasets that overlook tissue specificity and thus cannot anticipate off-target toxicity or resistance. Our work introduces a **context-aware, AI-driven framework** that integrates large-scale protein language model embeddings with tissue-resolved protein-protein interaction data to predict compatible and tissue-selective E3 ligases for any pathogenic target. By learning biochemical compatibilities directly from sequence space, this approach transcends the limitations of curated interaction databases and enables generalization to novel ligases or disease mutations. The resulting ligand–target rankings combine biochemical plausibility with spatial expression constraints, providing a scalable foundation for designing degraders that act precisely within disease-relevant tissues. This represents a conceptual and technical advance toward **precision degradomics** - a next generation of targeted protein degradation where therapeutic selectivity is defined not only by molecular affinity but also by cellular and tissue context. Subsequent steps will include structural integration for ternary complex modeling, biochemical validation of predicted pairs, and the deployment of a public platform to guide tissue-specific degrader discovery.

## 1. Background, motivation and problem statement

Over the past two decades, targeted protein degradation (TPD) has emerged as a powerful therapeutic strategy for tackling proteins once considered "undruggable," including drivers of cancer, inflammatory and autoimmune disorders, and neurodegenerative diseases [1]. TPD agents exploit the cell’s own ubiquitin-proteasome system (UPS): a bifunctional degrader first recruits an E3 ligase to its target and then facilitates poly-ubiquitin tagging, marking the protein for destruction [2]. The poly-ubiquitinated substrate is subsequently recognized by the 26S proteasome and processed into short peptides, free ubiquitin, and amino acids that the cell recycles, thereby eliminating the disease-associated protein and restoring proteostasis [3].

A key modality within TPD is the use of heterobifunctional molecules known as proteolysis-targeting chimeras (PROTACs), which simultaneously bind an E3 ubiquitin ligase and a target protein, bringing them into close proximity to facilitate ubiquitin transfer [4]. Ubiquitination is catalyzed through a highly conserved enzymatic cascade involving E1 activating enzymes, E2 conjugating enzymes, and E3 ligases [5]. E1 enzymes initiate the process by activating ubiquitin and transferring it to E2 enzymes, which are more diverse and act as intermediaries. E2 enzymes then collaborate with E3 ligases to transfer ubiquitin to specific substrate proteins [5]. Among these enzymes, E3 ligases are the most diverse and are critical for substrate specificity, as they determine which proteins are targeted for degradation [6,7,8,9]. As such, E3 ligases play a central role in UPS and are essential for the precision and effectiveness of targeted protein degradation strategies.

PROTACs are a novel class of small molecules with unique therapeutic potential and associated challenges [10]. PROTACs (Proteolysis-Targeting Chimeras) are heterobifunctional molecules designed to harness the cell’s ubiquitin-proteasome system by simultaneously binding a target protein and an E3 ubiquitin ligase, thereby forming a ternary complex that facilitates the ubiquitination and subsequent proteasomal degradation of the target [11]. This catalytic mechanism allows PROTACs to deplete disease-relevant proteins-including many considered undruggable by conventional small molecules-at sub stoichiometric doses and with sustained pharmacodynamic effects [2]. In contrast, Molecular Glues are monovalent compounds that enhance or stabilize interactions between an E3 ligase and a neo-substrate, redirecting endogenous degradation machinery toward new targets [12]. Although approximately 600 E3 ligases have been identified in the human proteome [13], PROTAC development has predominantly focused on just two-Cereblon (CRBN) and Von Hippel–Lindau (VHL) -due to their well-characterized structures and broad tissue expression [14].

However, reliance on this narrow subset of E3 ligases presents two major challenges. First, prolonged use of CRBN- or VHL-based degraders can impose evolutionary pressure, leading in the case of cancer to the emergence of resistance mutations or conformational changes that impair E3-ligase binding and reduce degradation efficacy [15].Second, the ubiquitous expression of CRBN and VHL across healthy and diseased tissues increases the risk of off-target protein degradation, raising concerns about systemic toxicity and limiting therapeutic selectivity [2]. While TPD offers a powerful mechanism to eliminate disease-driving proteins, its current implementations often lack tissue or cell-type specificity, undermining the principles of precision medicine [16]. Achieving true selectivity will require the development of degraders that engage context-specific E3 ligases or are activated conditionally—such as by tissue-specific expression, microenvironmental cues, or drug-induced control [17]. To address these limitations, precision-targeted protein degradation (TPD) strategies require the deployment of target- and tissue-enriched E3 ligases, which would enable the selective degradation of pathogenic proteins specifically within diseased cells or tissues [18]. This approach offers a promising path to minimize systemic toxicity, a major concern with current degraders that utilize ubiquitously expressed ligases such as CRBN and VHL [19]. By coupling substrate targeting E3 ligases with restricted expression patterns, degraders can be engineered to function only in relevant biological contexts, reducing off-target degradation in healthy tissues and enhancing therapeutic precision.

Expanding the use of tissue-enriched and tumor-selective E3 ligases could open new avenues for addressing diseases that have previously been intractable due to ligase incompatibility or systemic toxicity risks [20,21]. For example, ligases with expression enriched in the central nervous system, immune-cell subtypes, or tumor tissues—relative to healthy counterparts—could be leveraged to design degraders that act preferentially in those compartments, thereby minimizing off-target effects in non-diseased tissues [21]. Tissue- or context-specific E3 ligases may also offer unique structural motifs, distinct from commonly used ligases like CRBN and VHL, enabling the design of highly selective ligases and novel engagement mechanisms that further enhance the precision of targeted protein degradation [18,20]. A notable example of the challenges discussed above is CFT7455, a CRBN-recruiting PROTAC currently in clinical trials for Non-Hodgkin’s Lymphoma (NHL) and Multiple Myeloma (MM)—malignancies involving B cells and plasma cells, respectively [22] CFT7455 exerts its therapeutic effect by degrading the transcription factors IKZF1 (Ikaros) and IKZF3 (Aiolos), which are over-expressed or mutated in these hematologic cancers [23]. However, its systemic use also leads to the degradation of these targets in non-pathological cell types. This off-target activity manifests as dose-limiting toxicities—including neutropenia, peripheral neuropathy, immunosuppression, and broader hematopoietic suppression—observed in early clinical studies [24]. These adverse effects underscore the need for strategies that enable cell-type or tissue-restricted TPD when using E3 ligases with widespread expression.

Reducing the overreliance on a narrow subset of ubiquitously expressed E3 ligases and enhancing tissue specificity of targeted protein degradation (TPD) molecules could offer a strategic advantage in addressing current limitations [25]. While most clinically investigated PROTACs recruit ligases such as VHL or CRBN, these global E3 ligases are expressed across nearly all tissue types, increasing the risk of off-target degradation and dose-limiting toxicities due to their essential housekeeping functions [19]. Moreover, the widespread exposure of these ligases to degraders may accelerate the emergence of resistance mutations through selective evolutionary pressure [15].

E3 ligases exhibit tissue-enhanced or tissue-restricted expression profiles, offering an untapped opportunity to design tissue-selective degraders [13,18,26]. These localized E3 ligases often regulate context-specific protein networks or signaling pathways, and are less evolutionarily conserved, making them less susceptible to selective pressure and mutation-driven resistance. By leveraging these tissue-specific ligases, it may be possible to expand the functional E3 ligase toolbox, enabling more precise degradation of pathological proteins within disease-relevant tissues, while sparing non-target organs and reducing systemic toxicity. This approach aligns with the broader goal of advancing TPD as a precision medicine modality [25].

Hence, exploration of the broader E3 ligase landscape becomes imperative to address key challenges in targeted protein degradation, including off-target effects, acquired drug resistance, and the need for tissue-specific degradation [25]. The conventional reliance on a narrow subset of ubiquitously expressed ligases, such as CRBN and VHL, limits therapeutic precision and increases the risk of systemic toxicity [19]. These ligases, due to their broad expression and essential functions, are also subject to resistance mechanisms under prolonged selective pressure [15].

E3-ligase-substrate interaction (ESI) prediction has become the leading *in silico* strategy for probing the ubiquitin–proteasome landscape and prioritizing ligases for a chosen substrate or biological context. For example, UbiBrowser 1.0 [27] fused five evidence layers - domain enrichment, Gene Ontology overlap, co expression, phylogenetic similarity, and protein interaction topology - into a Bayesian network that outputs proteome scale ESI probabilities [27]. A new wave of deep learning approaches followed. E3 targetPred represents ligase and substrate sequences using composition of k spaced amino acid pairs (CKSAAP) to capture both local and long range dependencies before mapping them into a discriminative latent space for classification [28]. DeepUSI instead feeds one hot encoded full length protein sequences into a convolutional neural network that learns interaction motifs directly from raw sequence data, yielding state of the art accuracy on independent benchmarks [29]. ESI-NET, introduced by Chen et al. [30], is a multidimensional omics- and network-integrated framework that uses a Random-Forest model to identify E3 ubiquitin ligase-substrate interactions. UbiBrowser and ESI-NET are primarily feature-engineered frameworks that rely on curated biological and omics-derived descriptors, whereas DeepUSI and E3TargetPred learn E3–substrate interaction patterns directly from protein sequence using neural network-based models.

All four methods ultimately rely on curated repositories such as UbiBrowser annotations and BioGRID derived pairs. Hence, their predictive horizon is bounded by the completeness and quality of existing datasets. Existing methods are largely based on traditional machine learning approaches or early neural network architectures, which often struggle with overfitting and have limited capacity to model long-range dependencies in protein sequences. In contrast, protein language models have demonstrated a strong ability to capture long-range contextual relationships and have consistently achieved state-of-the-art performance across a range of protein engineering tasks. These properties make them a particularly suitable choice for E3– substrate interaction (ESI) prediction.

E3 ubiquitin ligases span a continuum of expression profiles in human tissues, ranging from ubiquitously expressed to tissue enhanced and truly tissue specific proteins [31]. Current computational ESI predictors - such as UbiBrowser, E3 targetPred and DeepUSI - have been trained on global interaction datasets compiled without tissue context, so the resulting scores are inherently tissue agnostic [27,29]. This constraint limits their capacity to address safety risks like off target toxicity or mutagenicity that can arise when degraders recruit ligases with broad physiological distribution [2]. Combining tissue resolved expression atlases with interaction evidence and harnessing the representational power of modern protein language models offers a biologically grounded path toward context aware ligase ranking that could help mitigate such liabilities [32,33,34].

For this purpose, we propose an AI-driven framework that predicts tissue-specific E3 ligases for pathogenically mutated target proteins. This approach is grounded in protein sequence data, with the model trained on billions of evolutionarily and functionally relevant sequence motifs. At its core is a protein language model that learns contextual embeddings capturing fine-grained sequence features—such as degrons, post-translational modification sites, and structural propensities—directly from raw amino acid sequences [35]. These high-dimensional embeddings are then used to assess the functional compatibility between E3 ligases and target proteins, integrating tissue specificity inferred from co-expression and enrichment patterns across transcriptomic and proteomic atlases.

Our model effectively ranks ligase–target pairs based on predicted binding compatibility, tissue-restricted expression, and the likelihood of forming productive ternary complexes. This enables mutation-selective degradation confined to disease-relevant tissues, while sparing essential proteins and avoiding off-target effects in healthy cells. Ultimately, this sequence-driven, transformer-enabled prediction platform represents a scalable and generalizable solution for rationally expanding the E3 ligase toolbox—unlocking context-specific degraders tailored to the molecular and spatial biology of disease.

## 2. Methods

Multiple protein language models (PLMs) have emerged as powerful tools for learning biologically meaningful representations directly from amino-acid sequences [37,38]. In this study, we evaluated three PLMs—ProtBERT, ESM-2 650M, and ESM-C 600M—which differ in model size, architecture, and, importantly, the scale and diversity of their pretraining data.

**ProtBERT** is a BERT-style encoder-only transformer with approximately **420 million parameters**, 30 transformer layers, and a 1,024-dimensional embedding representation. It was pretrained using masked-language modeling on approximately **216–217 million protein sequences from UniRef100** [36]. The use of UniRef100 exposes ProtBERT to a broad collection of protein sequences without the stronger sequence-identity reduction applied to UniRef50, providing substantial sequence diversity for learning transferable protein representations.

**ESM-2 650M** (esm2_t33_650M_UR50D) is a larger encoder-only transformer containing approximately 650 million parameters, 33 transformer layers, and a 1,280-dimensional embedding representation. It was pretrained on the **UniRef50-derived UR50/D 2021_04 dataset**, in which highly similar **sequences** are clustered to reduce redundancy [39,40]. Thus, compared with ProtBERT’s UniRef100 training corpus, ESM-2 emphasizes a more redundancy-reduced representation of protein sequence space.

**ESM-C 600M (ESM Cambrian)** represents a newer generation of protein language models and contains approximately **600 million parameters**, 36 transformer layers, and an embedding width of 1,152. Its pretraining dataset is substantially broader than those of ProtBERT and ESM- 2, combining sequences from UniRef, MGnify, and the Joint Genome Institute (JGI). After clustering at 70% sequence identity, these sources comprised approximately **8**3 million UniRef clusters, 372 million MGnify clusters, and 2 billion JGI clusters, respectively. ESM-C 600M was trained for 1.5 million steps on approximately 6.2 trillion amino-acid tokens. The inclusion of extensive metagenomic sequence data substantially expands the protein sequence diversity represented during pretraining [41].

The three PLMs therefore provide complementary sequence representations at different scales of pretraining: ProtBERT was trained on ∼217 million UniRef100 sequences; ESM-2 650M on the redundancy-reduced UniRef50/UR50D corpus; and ESM-C 600M on a much larger combination of UniRef, MGnify, and JGI data totaling approximately 2.46 billion sequence clusters and 6.2 trillion training tokens. Because the datasets are reported using different units—individual sequences for ProtBERT, a clustered UniRef corpus for ESM-2, and clusters plus training tokens for ESM-C—these quantities should not be interpreted as strictly equivalent measures of training-set size. Nevertheless, they illustrate the substantial expansion in sequence diversity and training scale represented by the newer PLMs.

All three models generate alignment-free, sequence-only representations, making them suitable for large-scale PPI prediction without requiring experimentally determined structures or multiple-sequence alignments [43]. As illustrated in Figure 1, two protein sequences are provided as input and independently encoded by the PLM. Their sequence-derived representations are subsequently combined into a joint representation and passed to the interaction-classification head, which produces a binary bound/unbound prediction together with an associated confidence score. Applying the same overall PPI-prediction framework to ProtBERT, ESM-2 650M, and ESM-C enables direct assessment of how differences in PLM architecture and pretraining scale affect ESI prediction.

**Figure 1.**
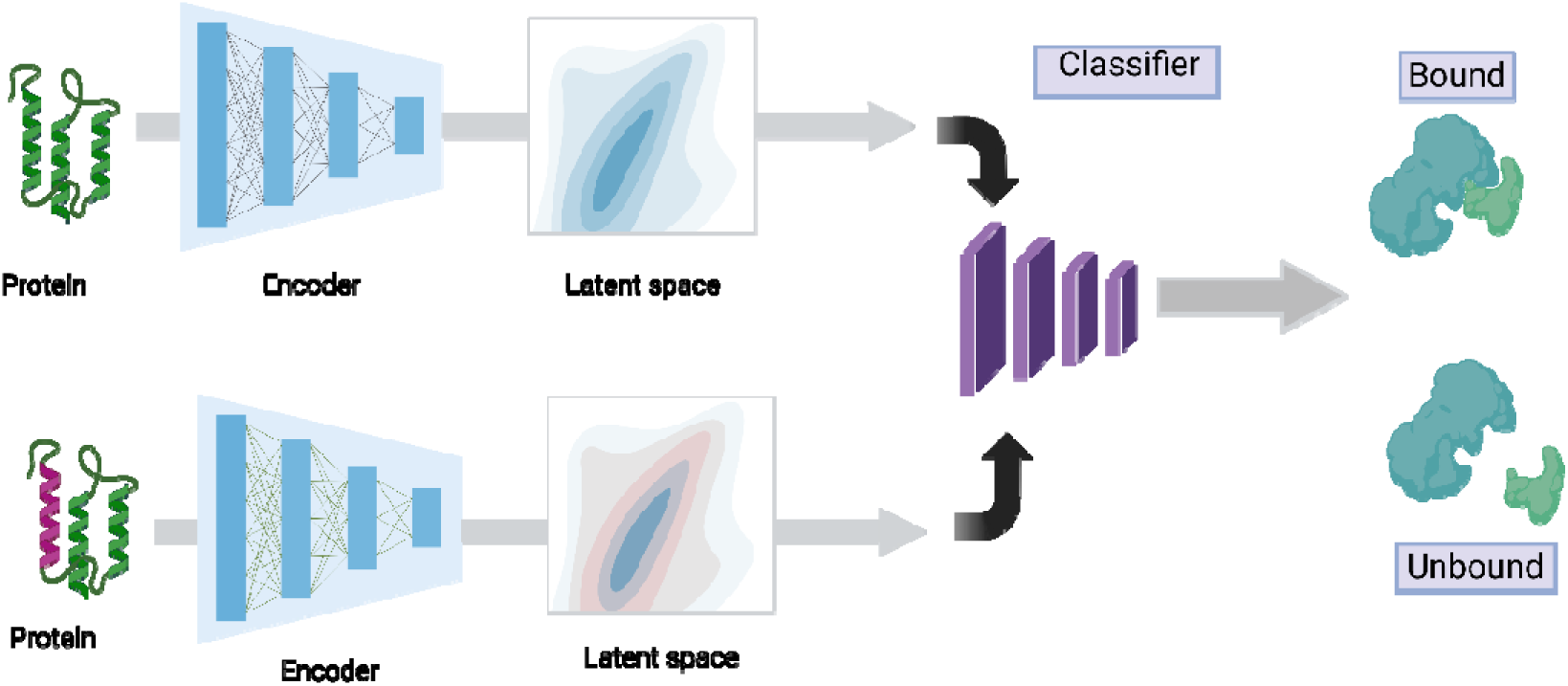
**– PLM based PPI Flow**

### 2.1. ELITE architecture

The classification head consists of only two linear layers a small head is streamlined for classification tasks with comparably small tissue specific datasets. There is a 20% dropout layer no activation function between the layers with a sigmoid final output function. The drop out value was determined empirically.

During training, raw protein sequences are processed by a context-aware encoder to produce high-dimensional residue embeddings. Paired embeddings are combined and passed through a two-layer classification head, with the full model optimized to distinguish interacting from non-interacting protein pairs. In the inference phase, both the encoder and classification head are fixed in their trained state: embeddings for unseen protein pairs are generated using the same encoder and then evaluated by the frozen classifier to yield interaction probabilities. The network training is driven by the optimization of the Binary Cross Entropy loss function.

Figure 2a provides a detailed view of the ELITE architecture. Each amino acid in the input protein sequence is converted into a dynamic embedding that captures long-range contextual dependencies, overcoming the limitations of static embeddings and the architectural constraints of RNN-based methods. A pooling layer condenses the sequence of residue embeddings into a single vector representation per protein, using a mechanism adapted from cross-domain architectures. The resulting protein embeddings are combined - typically via element-wise multiplication to generate a joint representation for the protein pair. This PPI embedding is then progressively reduced to a scalar interaction score through a series of dimensionality-reducing linear transformations. Figure 2b extends this architecture by explicitly incorporating tissue identity as a third model input through a learned tissue embedding, which is concatenated with the protein-pair representation before classification.

**Figure 2.**
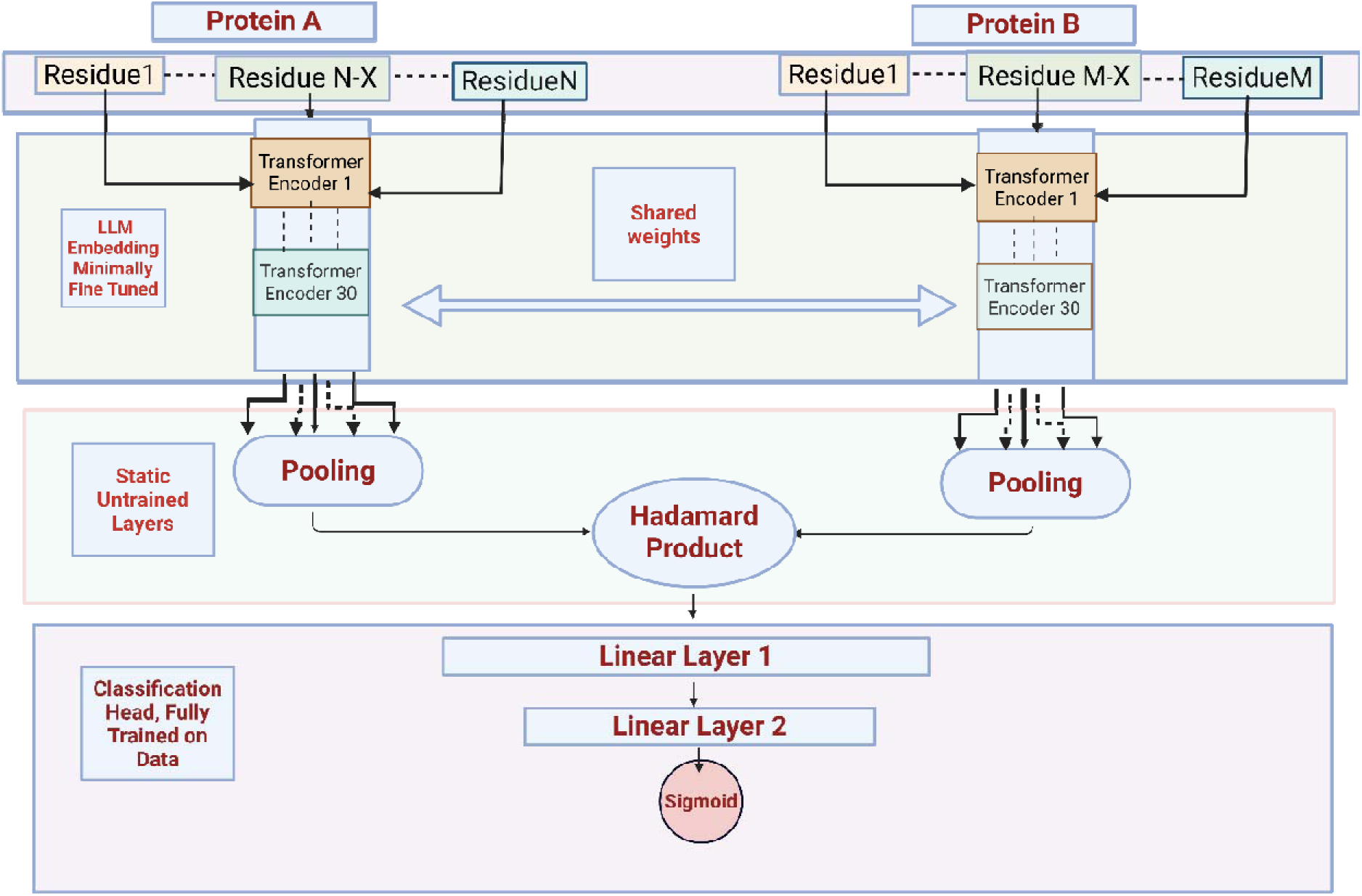
a: **Proposed ELITE architecture.**

**Figure 2b:**
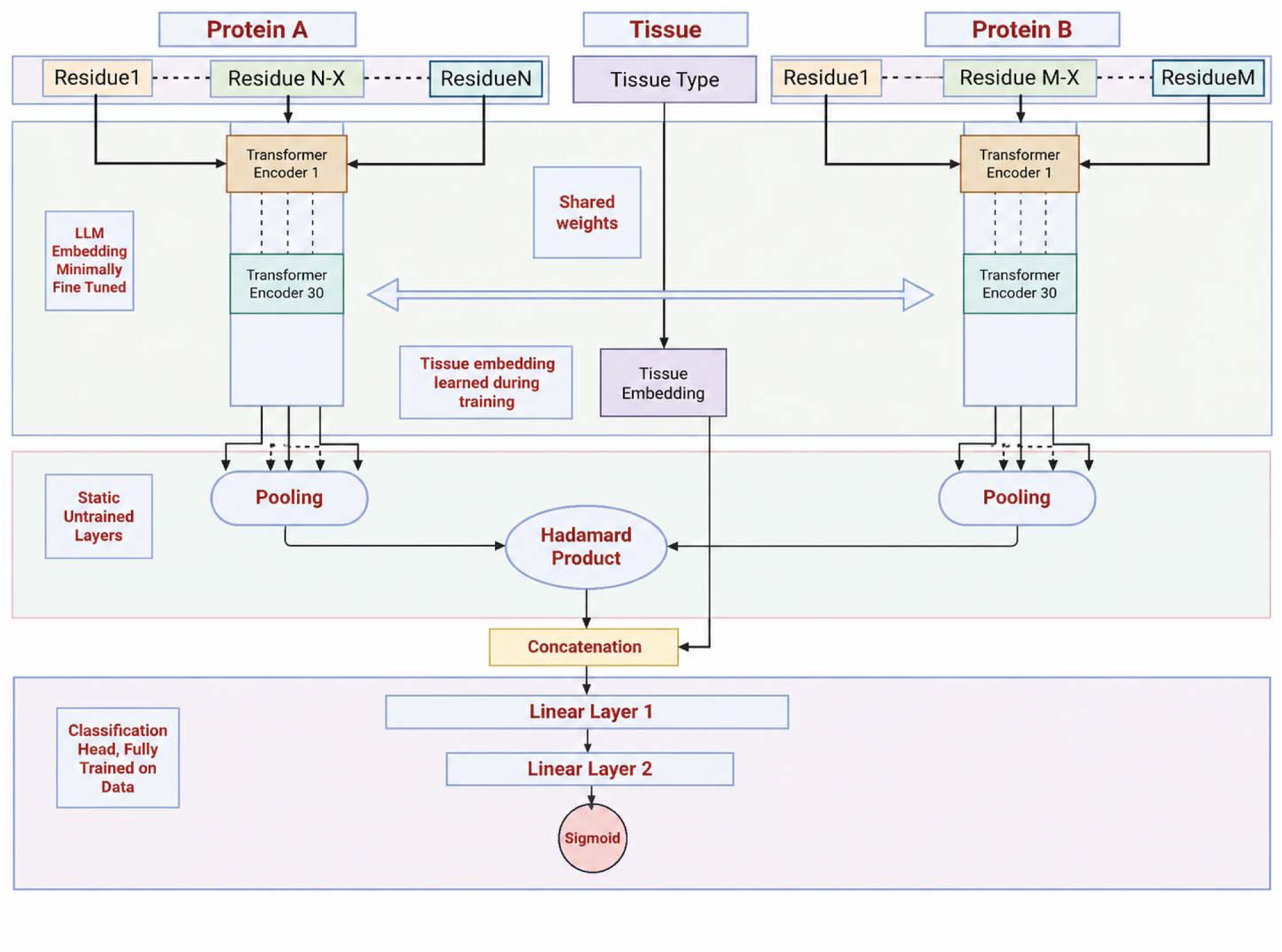
**ELITE architecture with tissue context as a third input. Protein A and Protein B are encoded by the shared-weight protein language model and pooled;**

### 2.2. Tissue-resolved protein-protein interactions

The Stanford Network Analysis Project (SNAP) is a comprehensive initiative that curates and maintains a diverse collection of network datasets spanning multiple domains, including social networks, communication systems, citation graphs, and collaboration networks [44]. Within SNAP, the BioSNAP subset serves as a specialized repository for biomedical network datasets, offering structured resources such as tissue-specific gene networks, drug-target interaction graphs, disease-gene associations, protein-protein interaction (PPI) networks, and more [45]. BioSNAP provides high-quality, well-documented datasets that support the development and benchmarking of computational methods in network biology, drug discovery, and biomedical data mining.

The tissue-specific PPI network utilized in our study was obtained from the BioSNAP collection. This network was originally constructed by the authors using a two-step approach involving the integration of tissue-specific gene expression profiles with a general PPI network to generate tissue-contextualized gene–gene associations [46]. We used this publicly available, pre-constructed network as the foundation for our analysis. The dataset obtained from the above procedure comprises a large number of small tissue-specific PPI networks (144 in total). To mitigate the partially small number of interactions per network, we first aggregated tissues into biologically meaningful groups, thereby reducing the number of datasets while enhancing the statistical and functional robustness of downstream analyses. We grouped the tissues based on criteria such as biological function, anatomical location, developmental origin, molecular signatures, and disease relevance, resulting in 26 consolidated tissue clusters (see Table S1 in Supplementary material for details). The ESIs were filtered by mapping the E3 Ligases as one of the interacting partners and removing duplicates. In the resulting networks, ESIs are treated as tissue-specific, reflecting the distinct regulatory environment within each tissue. However, the interacting entities - the E3 ligases and substrate proteins - may themselves exhibit broader expression patterns and are not necessarily confined to a single tissue. Dataset-level statistics are reported in Table 1.

**Table 1.** – Details of tissue specific E3 ligase - protein interaction data. The tuples in the last two columns indicate median, maximum and minimum of the number of amino acids of proteins.

| SI No. | Name | No. of ESI | No. of E3 Ligases | No. of Substrates | E3 Ligase Size Distribution | Substrate Size Distribution |
| --- | --- | --- | --- | --- | --- | --- |
| 1 | ALL Tissues | 10,498 | 299 | 345 | (508,34350,70) | (542,5202,84) |
| 2 | Cardiac | 522 | 98 | 419 | (510,34350,70) | (604,4911,84) |
| 3 | Adrenal | 364 | 82 | 301 | (521,3159,95) | (557,4911,84) |
| 4 | Bone | 420 | 83 | 347 | (509,3159,95) | (568,4911,84) |
| 5 | Brain | 541 | 97 | 437 | (510,4646,70) | (574,4911,84) |
| 6 | Renal | 223 | 101 | 429 | (515,5202,70) | (574,4911,84) |
| 7 | PNS | 187 | 63 | 165 | (510,2843,105) | (574,4374,84) |
| 8 | Immune | 502 | 93 | 405 | (509,4646,70) | (574,4911,84) |
| 9 | Auditory | 25 | 10 | 25 | (464,2414,113) | (875,1512,309) |
| 10 | Blood | 494 | 90 | 402 | (508, 4646, 70) | (574,4911,84) |
| 11 | Reproductive Female | 453 | 90 | 367 | (520,5202,70) | (574,4911,84) |
| 12 | Mammary | 298 | 59 | 253 | (524,3159,101) | (568,4911, 303) |
| 13 | Endocrine | 377 | 83 | 311 | (515,3159,95) | (568,4911, 84) |
| 14 | Fetal | 531 | 94 | 431 | (508,5202,70) | (592, 4911, 84) |
| 15 | Gastrointestinal | 540 | 101 | 436 | (510,5202,70) | (604,4911,84) |
| 16 | Hepatic | 164 | 60 | 142 | (531,2843,124) | (562,4374,84) |
| 17 | Oral | 334 | 79 | 279 | (510,5202,95) | (568,4374, 84) |
| 18 | Pulmonary | 531 | 99 | 419 | (510,5202,70) | (592,4911, 84) |
| 19 | Reproductive Male | 501 | 94 | 403 | (510, 5202,70) | (574,4911,84) |
| 20 | Skeletal Muscle | 476 | 92 | 381 | (513,4646,70) | (570,4911,84) |
| 21 | Skin | 404 | 84 | 333 | (510,3159,95) | (568,4911,84) |
| 22 | Spinal Cord | 542 | 97 | 437 | (510,4646,70) | (574,4911,84) |
| 23 | Thyroid | 373 | 81 | 301 | (520,5202,95) | (568,4911,84) |
| 24 | Urinary Tract | 237 | 50 | 205 | (511,5202,101) | (568,4911,309) |
| 25 | Vascular | 352 | 79 | 295 | (510,2843,95) | (570, 4374,84) |
| 26 | Vision | 378 | 84 | 311 | (510,5202,95) | (568,4374,84) |
| 27 | Connective | 399 | 86 | 322 | (570,4911,84) | (510,3159,95) |

As part of the data-preparation workflow for machine learning, the tissue-specific gene-gene interaction networks were converted into binary PPI networks using Entrez-to-UniProt ID mappings. Because a single gene can produce multiple protein isoforms, one gene–gene interaction can correspond to several possible protein–protein interactions (PPIs). Many of these PPIs are therefore only predicted from the possible combinations of isoforms and have not been directly validated experimentally. A subsequent filtering step described below removes biologically implausible or weakly supported interactions, improving data confidence.

To identify experimentally supported interactions within the hypothetical tissue-specific gene networks, it is necessary to compare them against curated PPI datasets derived from experimental studies. Several well-established PPI repositories are available for this purpose, including the Biological General Repository for Interaction Datasets [47,48], Search Tool for the Retrieval of Interacting Genes/Proteins (STRING) [49,50], IntAct [51], Molecular INTeraction Database (MINT) [52], and the Human Protein Reference Database (HPRD). Each of these databases contains hundreds of thousands of PPIs but differs in data format, confidence in scoring systems, curation standards, and completeness. Integrating these heterogeneous datasets poses significant challenges due to the lack of standardized data models, inconsistent identifiers, and contradictory interaction evidence arising from methodological differences and varying experimental conditions. The integration process is therefore resource-intensive and falls outside the scope of this study. As an alternative, we explored the use of pre-integrated and standardized PPI datasets, which offer harmonized formats, unified confidence scoring, and broader interaction coverage-providing a practical solution for mapping hypothetical interactions to experimentally validated protein-protein interactions.

The Agile Protein Interaction DataAnalyzer (APID) is a comprehensive and high-quality PPI database constructed by integrating data from several widely used repositories, including BIND [53], DIP [54], HPRD , IntAct [51], and MINT[52]. APID adheres to the Proteomics Standards Initiative – Molecular Interaction (PSI-MI) format [55], incorporating detailed metadata such as interactor information, interaction evidence, PubMed references (PMIDs), confidence scores, biological roles, and interaction types (e.g., direct, indirect, complex). Experimental evidence in APID is derived from multiple methods: high-throughput techniques such as yeast two-hybrid (Y2H) and affinity-purification mass spectrometry (AP-MS) contribute 50-60% of entries, biochemical methods like co-immunoprecipitation (Co-IP) and pull-down assays account for 20- 25%, structural and high-resolution techniques for 10-15%, and fluorescence-based assays and biophysical methods (e.g., single-molecule microscopy, isothermal titration calorimetry) each provide 5-10% of the total entries [56]. The APID database is structured into three quality tiers: Levels 0-2. APID Level 2 (the highest level) was chosen for further filtering. This Level includes 22,707 proteins and 154,955 interactions [56]. This curated, experimentally validated interaction dataset serves as the primary source for downstream transformation, filtering, and modeling.

To assemble the training corpus, we intersected each tissue-specific gene network with APID Level 2, yielding 26 tissue-resolved ESI datasets that constituted the positive class. For the negative class, we adopted the rigorously curated Negatome 2.0 non-interaction set released by Hamp and Rost, which minimizes label noise by excluding protein pairs with any experimental or structural evidence of interaction [57]. All interacting partners were mapped to canonical sequences via the UniProt Knowledgebase [58], producing tissue-specific sequence pairs labeled as interacting or non-interacting. These pairs formed the input to our machine learning pipeline. Figure 3 summarizes the complete data-engineering workflow and introduces the modeling architecture described in subsequent sections.

**Figure 3.**
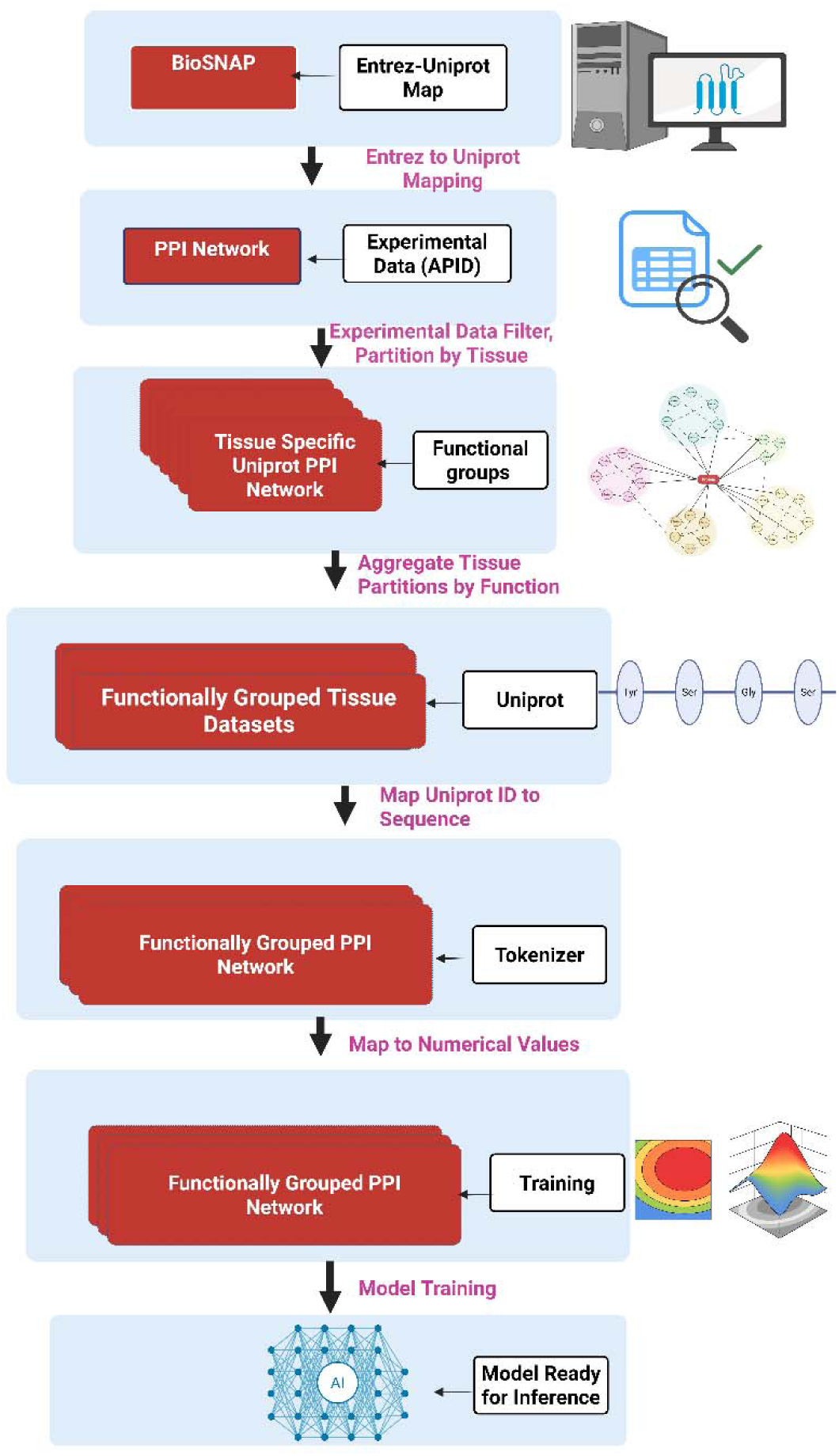
**- Data Engineering and ML Flow**

### 2.3. Model training and evaluation

Prior ESI predictors are typically trained from scratch on engineered datasets. ELITE follows the same overarching objective while fine-tuning a pretrained PLM backbone with a classification head on tissue-specific datasets. For evaluation, we report AUPR and AUROC, aligning with common practice to ensure consistency with the literature and robust, comparable performance assessments.

We report both aggregate and point-wise metrics to gauge the model’s ability to detect positives, whether common or rare. In imbalanced data such as ours, Precision-Recall Area Under the Curve (PR-AUC), precision, and recall are especially informative, because together they impose tight control over false positives and false negatives across decision thresholds [59,112]. By contrast, a model can attain a high Receiver Operating Characteristics Area Under the curve (ROC-AUC) even while producing many false positives or false negatives, so ROC-AUC alone would overstate performance [60,112].

We evaluated the model using **stratified 10-fold cross-validation**. The full dataset was divided into ten equal-sized folds that each preserved the original class distribution. In every iteration, nine folds were used for training while the remaining fold served as the validation set, ensuring that every sample was tested exactly once. This approach maintains class balance within each split, reduces variance from sampling error, and yields a more reliable and less biased estimate of real-world performance than a single hold-out split or non-stratified k-fold validation [61,62,63,64,65]. Specific training details are provided in the Supplementary Material Training section.

### 2.4. Making model predictions explainable

Integrated Gradients (IG) [66,67,68] attributes a model’s output F(x) to each feature by integrating the gradient along the straight path from a baseline (neutral) to the input. The mathematical formulae is shown in <u>Eq 1.</u>

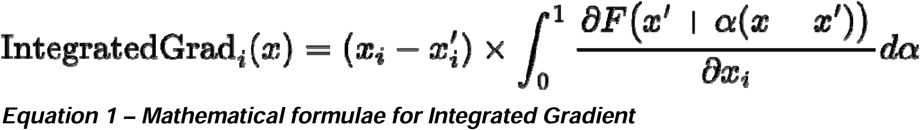

Integrated Gradients (IG) was used to elucidate which protein domains and individual residues most strongly drive our BERT-based PPI predictions. IG’s proven success on biological sequence models provides robust validation for high-confidence interaction calls [69]. Crucially, IG adheres to its two foundational axioms in a biologically meaningful way: it assigns nonzero attributions to any residue whose isolated perturbation changes the interaction score-thereby pinpointing known "hot-spot" amino acids (Sensitivity)-and it produces identical attribution maps across different transformer architectures that yield the same PPI outputs, whether ProtBERT, ESM-1b, or a custom fusion head is used (Implementation Invariance).

#### 2.4.1. Implementation notes

Integrated Gradients "steps" (n_steps) is how many intermediate points you sample as you move smoothly from a baseline input to the actual input to approximate the integral of gradients along that path: IG evaluates mixed inputs of the form baseline + α·(input−baseline) for α values between 0 and 1, computes gradients at each point, averages/sums them, and scales by (input−baseline) to produce attributions; in this implementation it was done with the Captum library using 50 steps, where more steps usually gives a smoother, more accurate estimate but costs more compute. Here, the baseline is an all-[PAD] reference, which works well because PAD represents "no residue" while still being a valid, same-shaped input; with the attention mask, padded positions are meant to be ignored by the model, so attributing changes from PAD to real residues cleanly reflects which residue content drives the PPI prediction.

## 3. Results and use cases

### 3.1. Cross-validated prediction performance

Tables <u>2A</u> and <u>2B</u> summarize the ten-fold stratified cross-validation PR-AUC and ROC-AUC results across tissues for ELITE, ESM-2 650M, and ESM-C 600M. The model attains consistently high ROC-AUC and PR-AUC scores, demonstrating not only strong class separation but also reliable prioritization of the positive class-i.e., low false-positive and false-negative rates. The sole outlier is the auditory tissue, where ROC-AUC is poor. This is attributable to an insufficiently small training set that cannot adequately supervise fine-tuning. However, even in smaller datasets like the one discussed, the model reports a high PR-AUC showing a high precision for the small number of positive predictions.

Hepatic tissue also shows comparatively modest ROC-AUC values, again reflecting limited sample sizes. Overall, a trend emerges: larger tissue-specific datasets yield proportionally higher metrics, underscoring the model’s capacity to scale rapidly with data volume and approach maximal performance.

### 3.2. Pooled all-tissue evaluation with tissue embedding

To evaluate performance across heterogeneous biological contexts while retaining tissue identity, all 26 tissue-specific E3-substrate interaction datasets were combined into a single pooled dataset, with tissue retained as an explicit model feature. The pooled dataset comprised 299 unique E3 ligases and 345 unique substrates. The pooled proteins spanned 70–34,350 amino acids for E3 ligases (median 508) and 84–5,202 amino acids for substrates (median 542). For each protein pair, the tissue label was mapped to a learned tissue embedding and combined with the pooled protein-pair representation before the prediction head, as illustrated in Figure 2b. This pooled tissue-aware evaluation was performed only with ESM-C 600M using 10 × 10 cross-validation and achieved a ROC-AUC of 99.3% and a PR-AUC of 99.9%. ProtBERT and ESM-2650M were not evaluated in this pooled setting as they were already evaluated on 26 tissues. These results indicate that ESM-C retains near-ceiling discrimination and positive-class prioritization when interactions from all tissues are modeled jointly while tissue context is explicitly represented.

### 3.3. Testing under balanced positive-to-negative ratio

ProtBERT was evaluated under both balanced (1:1) and imbalanced (1:9) positive-to-negative ratios to assess the stability of its predictive performance across different class distributions. Consistent superiority under both settings would indicate that the model’s advantage is not restricted to a single sampling regime, while the 1:9 evaluation provides a more realistic benchmark for highly imbalanced E3-substrate interaction prediction.

Comparing the two class-ratio settings, the overall pattern suggests that the model is **broadly robust**, with most tissues maintaining good predictive performance regardless of whether the evaluation is balanced (**1:1**) or imbalanced (**9:1**). In the **1:1** setting, many tissues such as **Adrenal, Bone, Reproductive Female, Mammary, Oral, Skin, Spinal Cord, Thyroid, Urinary Tract, Vascular, and Vision** show particularly strong results, with high PR-AUC, precision, recall, and accuracy. In the **9:1** setting, most tissues remain similarly strong, again showing high PR-AUC and recall with only modest variation in precision and accuracy.

Detailed balanced-ratio results are provided in Supplementary Table S2.

### 3.4. Benchmark against alternative methods

Among the existing computational approaches for ESI prediction, four representative methods include UbiBrowser, ESI-NET, DeepUSI, and E3TargetPred. UbiBrowser and ESI-NET are feature-engineered frameworks built on extensive biological and omics-derived data integration, which are not straightforward to reproduce or re-implement. In contrast, DeepUSI and E3TargetPred are sequence-based deep learning models that infer interaction patterns directly from protein sequences. We therefore use DeepUSI and E3TargetPred as direct sequence-based method benchmarks, while ESM-2 650M and ESM-C 600M are included as alternative protein-language-model backbone comparisons under the same tissue-aware prediction setting. This design enables comparison both against established E3-substrate prediction architectures and against newer large-scale protein foundation models.

We used PR-AUC as the primary performance metric because of the high-class imbalance in the dataset. The large number of true negatives can make ROC-AUC appear deceptively strong, as small changes in false positives have limited effect on the false-positive rate [112]. E3TargetPred was executed from the public codebase using the repository’s initial configuration on a single stratified 90:10 train:test split (positive:negative = 1:9), and DeepUSI was included as an additional published deep learning baseline. E3TargetPred uses CKSAAP sequence features for each protein and learns a joint autoencoder-plus-classifier in which the latent embedding must both reconstruct a pair feature vector from the two singles and distinguish true E3-target pairs from non-pairs [28]. Three key hyperparameters shape its behavior: *k* (the CKSAAP gap) controls sequence-context granularity, *LV* (latent dimensionality) determines embedding capacity and interpretability, and λ balances reconstruction fidelity against discriminative separation, trading expressive detail for class compactness in the latent space. By contrast, DeepUSI takes raw amino acid sequences as input and uses a convolutional neural network pipeline in which each sequence is encoded through an embedding layer, followed by three convolution operations, global max pooling, and a fully connected layer; the resulting protein embeddings are then combined and passed through a dropout-regularized neural network to generate the final E3–substrate interaction score [29].

As shown in Table 3, ESM-C 600M provides the strongest overall performance among the evaluated models. Across the 26 tissues, ESM-C achieves a mean AUPR of 99.24%, compared with 97.73% for ELITE/ProtBERT and 83.82% for ESM-2 650M. ESM-C records the highest tissue-level AUPR in 25 of 26 tissues; the only exception is the auditory dataset, where ELITE/ProtBERT (89.00%) is marginally higher than ESM-C (88.87%). The advantage of ESM- C is also evident relative to the re-implemented sequence-based baselines: DeepUSI averages 91.54% AUPR and E3TargetPred 70.79% across the 24 tissues for which their results are available. Thus, ESM-C is the best-performing model in the benchmark overall, whereas ELITE remains highly competitive and substantially outperforms ESM-2 650M, DeepUSI, and E3TargetPred on mean AUPR. These results suggest that the broader and more recent pretraining regime of ESM-C provides representations that transfer particularly effectively to tissue-resolved E3-substrate interaction prediction.

**Table 2.** A - **PR-AUC results of 10-fold cross-validation across 26 tissue-specific datasets, with an additional pooled ALL Tissues evaluation (serial 27).**

| Sl. | Tissue | ESM-C 600M | ELITE / ProtBERT | ESM-2 650M |
| --- | --- | --- | --- | --- |
| 1 | Cardiac | 99.64 | 98.00 | 87.63 |
| 2 | Adrenal | 99.71 | 99.00 | 87.23 |
| 3 | Bone | 99.77 | 98.00 | 87.19 |
| 4 | Brain | 99.72 | 98.00 | 87.72 |
| 5 | Renal | 99.74 | 98.00 | 84.00 |
| 6 | PNS | 99.25 | 97.00 | 85.58 |
| 7 | Immune | 99.66 | 98.00 | 86.41 |
| 8 | Auditory | 88.87 | 89.00 | 9.61 |
| 9 | Blood | 99.67 | 98.00 | 85.70 |
| 10 | Reproductive F | 99.70 | 98.00 | 89.28 |
| 11 | Mammary | 99.55 | 99.00 | 86.16 |
| 12 | Connective | 99.69 | 99.00 | 86.95 |
| 13 | Endocrine | 99.75 | 99.00 | 87.50 |
| 14 | Fetal | 99.73 | 98.00 | 86.17 |
| 15 | Gastrointestinal | 99.70 | 98.00 | 84.69 |
| 16 | Hepatic | 99.07 | 97.00 | 81.81 |
| 17 | Oral | 99.59 | 98.00 | 88.06 |
| 18 | Pulmonary | 99.73 | 96.00 | 85.54 |
| 19 | Reproductive M | 99.89 | 98.00 | 87.06 |
| 20 | Skeletal | 99.71 | 98.00 | 85.43 |
| 21 | Skin | 99.53 | 99.00 | 89.16 |
| 22 | Spinal Cord | 99.76 | 98.00 | 87.33 |
| 23 | Thyroid | 99.65 | 99.00 | 88.28 |
| 24 | Urinary Tract | 99.51 | 97.00 | 87.42 |
| 25 | Vascular | 99.64 | 99.00 | 87.06 |
| 26 | Vision | 99.88 | 98.00 | 90.44 |
| 27 | ALL Tissues | 99.90 | - | - |

**Table 2.** B - ROC-AUC results of 10-fold cross-validation across 26 tissue-specific datasets, with an additional pooled ALL Tissues evaluation (serial 27). Values are shown as percentages to two decimal places.

| Sl. | Tissue | ESM-C 600M | ESM-2 650M | ELITE / ProtBERT |
| --- | --- | --- | --- | --- |
| 1 | Cardiac | 98.42 | 96.15 | 92.00 |
| 2 | Adrenal | 98.63 | 95.45 | 93.00 |
| 3 | Bone | 98.80 | 95.56 | 93.00 |
| 4 | Brain | 98.43 | 95.82 | 91.00 |
| 5 | Renal | 98.54 | 94.04 | 90.00 |
| 6 | PNS | 96.66 | 93.44 | 90.00 |
| 7 | Immune | 98.23 | 95.39 | 90.00 |
| 8 | Auditory | 41.93 | 36.27 | 49.00 |
| 9 | Blood | 98.51 | 94.64 | 92.00 |
| 10 | Reproductive F | 98.45 | 96.89 | 94.00 |
| 11 | Mammary | 97.87 | 94.94 | 96.00 |
| 12 | Connective | 98.34 | 95.72 | 95.00 |
| 13 | Endocrine | 98.81 | 95.52 | 95.00 |
| 14 | Fetal | 98.62 | 95.48 | 93.00 |
| 15 | Gastrointestinal | 98.28 | 94.75 | 91.00 |
| 16 | Hepatic | 96.12 | 92.19 | 81.00 |
| 17 | Oral | 98.03 | 95.73 | 95.00 |
| 18 | Pulmonary | 98.76 | 95.03 | 85.00 |
| 19 | Reproductive M | 99.31 | 95.58 | 94.00 |
| 20 | Skeletal | 98.55 | 94.93 | 91.00 |
| 21 | Skin | 97.88 | 96.17 | 95.00 |
| 22 | Spinal Cord | 98.67 | 95.66 | 92.00 |
| 23 | Thyroid | 98.30 | 96.00 | 96.00 |
| 24 | Urinary Tract | 98.07 | 95.26 | 92.00 |
| 25 | Vascular | 98.24 | 95.24 | 95.00 |
| 26 | Vision | 99.25 | 96.75 | 94.00 |
| 27 | ALL Tissues | 99.30 | - | - |

**Table 3:**
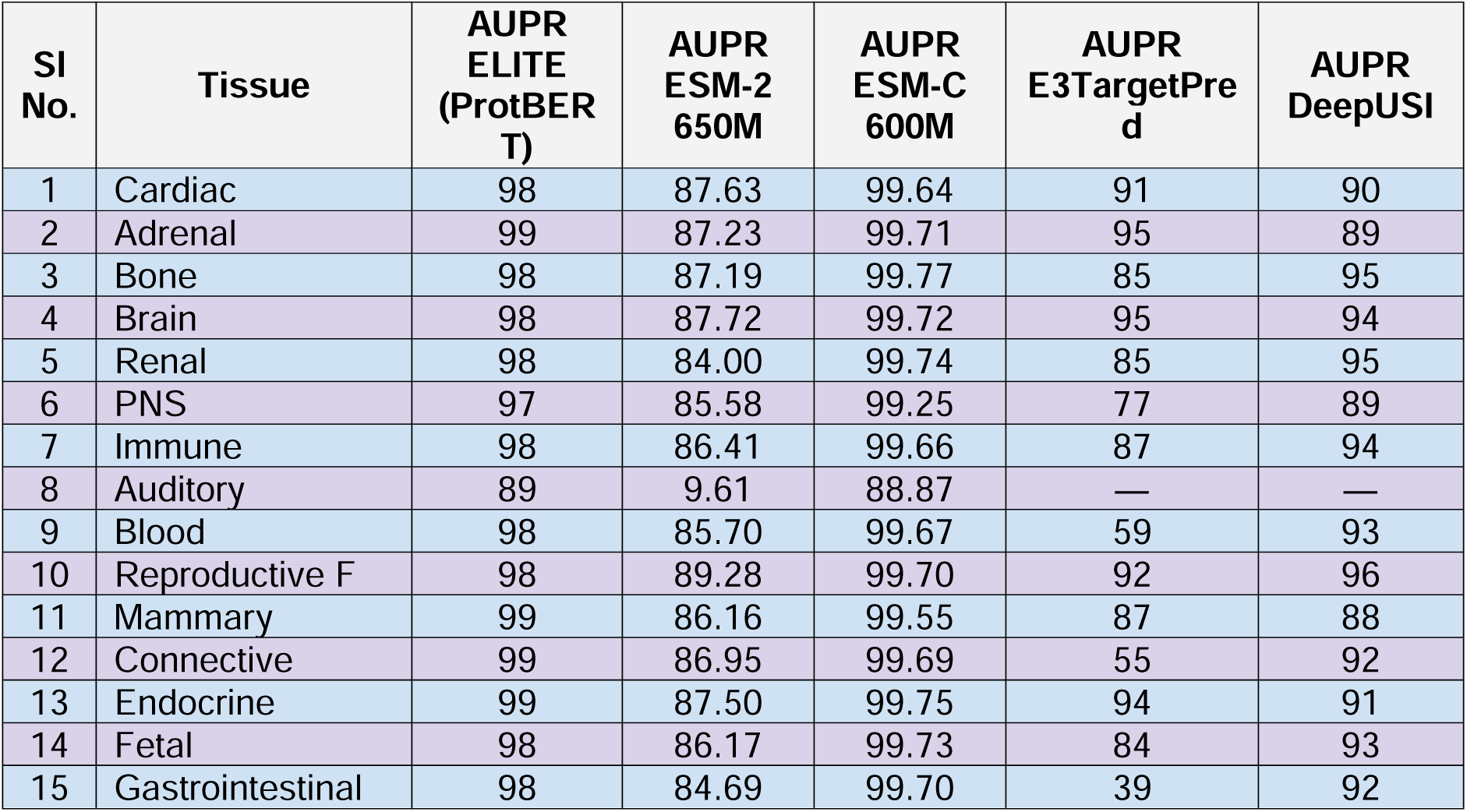
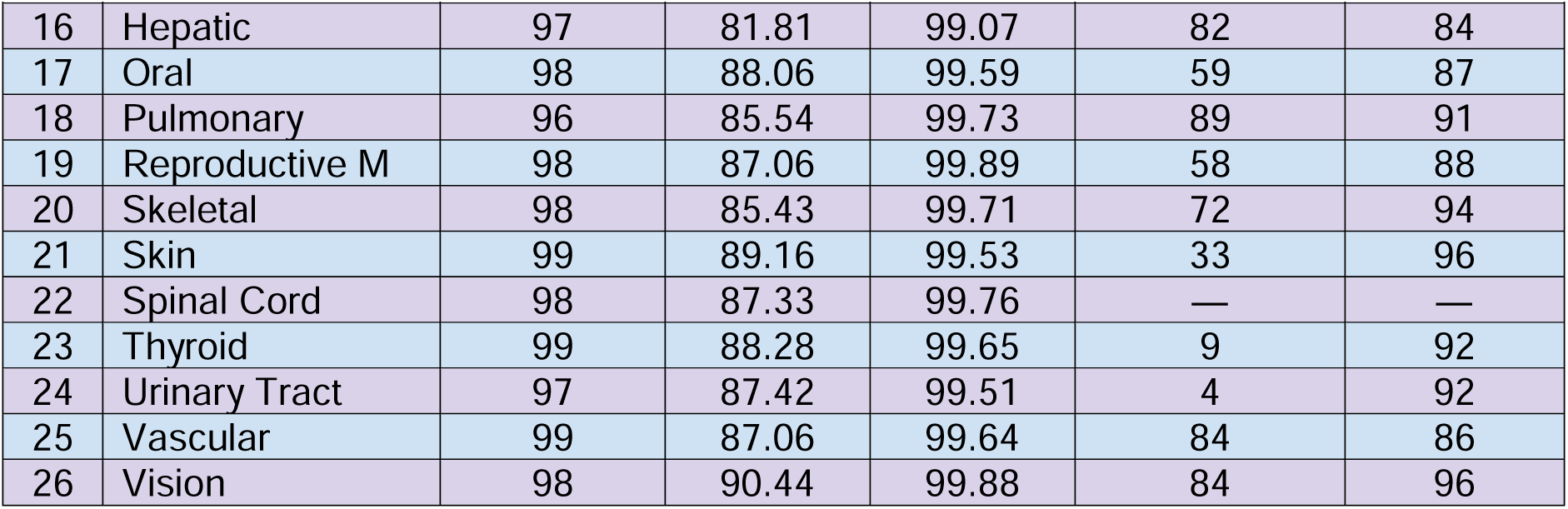
Tissue-wise AUPR comparison of ELITE (ProtBERT), ESM-2 650M, ESM-C 600M, E3TargetPred, and DeepUSI. Values are percentages; dashes indicate unavailable baseline results.

### 3.5. Example application 1: Targeting an intractable pathogenic transcription factor in gastrointestinal cancers

Transcription factors (TFs) are challenging targets. Wild-type p53 regulates the expression of >500 genes controlling cell-cycle arrest, DNA repair, and apoptosis, and its loss-of-function mutations drive many cancers. Likewise, NF-κB governs transcription of 500-1000 genes implicated in inflammation, immune evasion and tumor progression [70]. Despite their central roles, classical TFs lack accessible ligand-binding pockets, and no FDA-approved therapies yet directly disrupt protein-DNA interactions of p53, NF-κB, or similar TFs.

Despite their central role in disease-relevant pathways, transcription factors have historically been considered undruggable due to their lack of deep binding pockets, high structural flexibility, and primarily nuclear localization [71,72]. While small molecules have successfully targeted members of the nuclear receptor superfamily (a structurally distinct subset of TFs with ligand-binding domains), no FDA-approved therapies directly inhibit classical transcription factors like p53 or NF-κB at the protein-DNA interaction level [73,74].

Mutated p53 is one of the most extensively studied targets in cancer biology, owing to its pivotal tumor-suppressive functions and its status as the most frequently mutated gene across human cancers [75,76]. In its wild-type form, p53 is ubiquitously expressed and plays a critical role in maintaining genomic stability. Upon cellular stress or DNA damage, p53 activates the transcription of genes involved in DNA repair; if repair mechanisms fail, it initiates apoptosis through the induction of pro-apoptotic genes [76]. Over the past five decades, p53 has been the focus of intense research, leading to the identification of 3124 distinct variants [75]. These variants are classified based on predicted clinical significance into categories such as pathogenic, likely pathogenic, benign, likely benign, or of uncertain significance.

In this study, we focus on a specific p53 variant (R175H), which not only loses its native tumor-suppressive function but also acquires novel oncogenic properties that contribute to increased tumor aggressiveness [77]. Such gain-of-function mutations in p53 represent promising targets for therapeutic intervention, particularly through targeted protein-degradation strategies. Accordingly, we propose hypotheses regarding tissue-specific E3 ligases that are predicted to exhibit high binding affinity toward the mutant p53 protein, thereby enabling selective degradation. Gastrointestinal (GI) cancers - encompassing malignancies of the esophagus, stomach, small and large intestines, liver, and pancreas—represent some of the most lethal cancer types. Collectively, GI cancers account for approximately 26% of all cancer diagnoses and contribute to 35-90% of cancer-related mortality, underscoring the urgent need for effective, precision-targeted therapies [78]. The ranked p53 R175H E3-ligase predictions and corresponding ligase annotations are presented in Table 4 and Table 5.

**Table 4.** - Predictions - Ranked E3Ls in p53 mutation in GI Cancer.

| SI No. | E3 Ligase Uniprot ID | Score | RNA Seq HPA | RNA Seq GTEx | FANTOM | Sc RNA Cell Type |
| --- | --- | --- | --- | --- | --- | --- |
| 1 | Q9UKV5 | 0.99 | GI | GI | GI | GI |
| 2 | O00623 | 0.99 | GI | Non-Specific | Non-Specific | Non-Specific |
| 3 | Q5T447 | 0.98 | GI | GI | GI | GI |
| 4 | O00237 | 0.93 | GI | GI | GI | GI |
| 5 | Q86UW9 | 0.90 | GI | GI | GI | GI |
| 6 | Q9ULV8 | 0.90 | GI | GI | GI | GI |

**Table 5.** – Details of Ligases in Table.

| SI No. | E3 Ligase Uniprot ID | Entrez Gene ID | Official Name |
| --- | --- | --- | --- |
| 1 | Q9UKV5 | AMFR | E3 ubiquitin-protein ligase AMFR |
| 2 | O00623 | PEX12 | Peroxisome assembly protein 12 |
| 3 | Q5T447 | HECTD3 | E3 ubiquitin-protein ligase HECTD3 |
| 4 | O00237 | RNF103 | E3 ubiquitin-protein ligase RNF103 |
| 5 | Q86UW9 | DTX2 | Probable E3 ubiquitin-protein ligase DTX2 |
| 6 | Q9ULV8 | CBLC | E3 ubiquitin-protein ligase CBL-C |

Figures 4a, <u>4b</u>, <u>4c</u>, <u>4d</u>, <u>5a</u>, and <u>5b</u> map residue-level attribution scores for the interactions between the E3 ligases RNF103 and DTX2 and the cancer-associated p53 R175H variant, with red and green intensities marking residues that respectively promote or hinder binding. Both ligases show strong reliance on p53’s intrinsically disordered, proline-rich stretch spanning residues 60 – 85 (Figure 4 <u>c</u>), a region known to interact with HRMT1L2, WWOX, and CCAR2 [33,79,80] - whereas the structured β-strand at residues 250-270 (Figure 4 <u>d</u>), which normally accommodates partners such as HIPK1, ZNF385A, and AXIN1 [81,82,83] shows strongly negative attribution. This reflects a general principle: intrinsically disordered regions (IDRs) enable multivalent, adaptive interactions, while structured domains impose spatial and chemical constraints that may favor or disfavor specific partners depending on compatibility [84].

**Figure 4.**
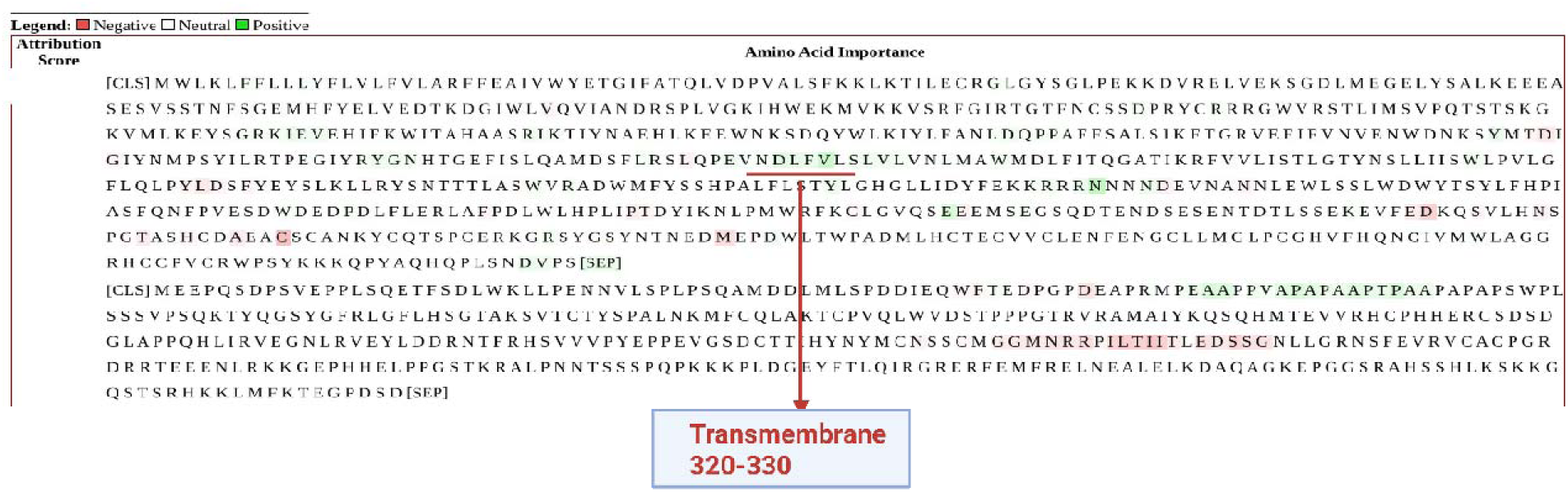
a – **O00237 (RNF103) transmembrane segment and p53 R175H interaction**

**Figure 4b.**
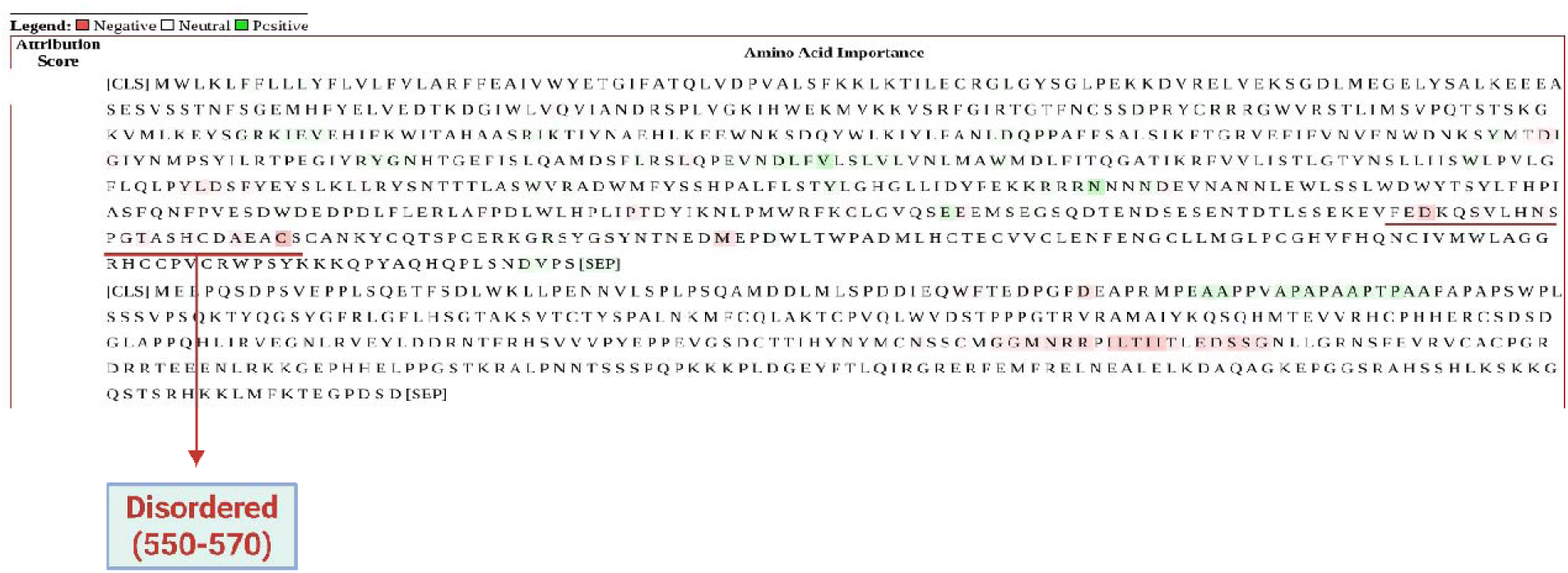
**- Disordered region in RNF103 (O00237) linked to p53 R175H**

**Figure 4c.**
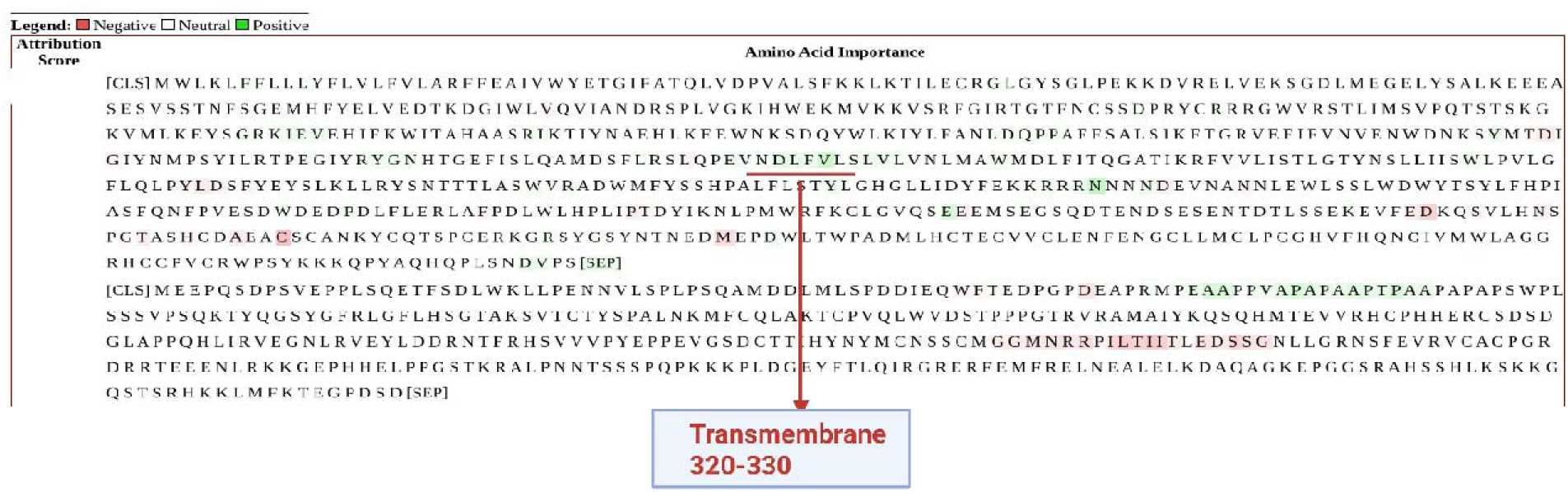
**- Pro-rich disordered region (60–85); interactors: WWOX, HRMT1L2, CCAR2; RNF103 (O00237) linked to p53 R175H**

**Figure 4d.**
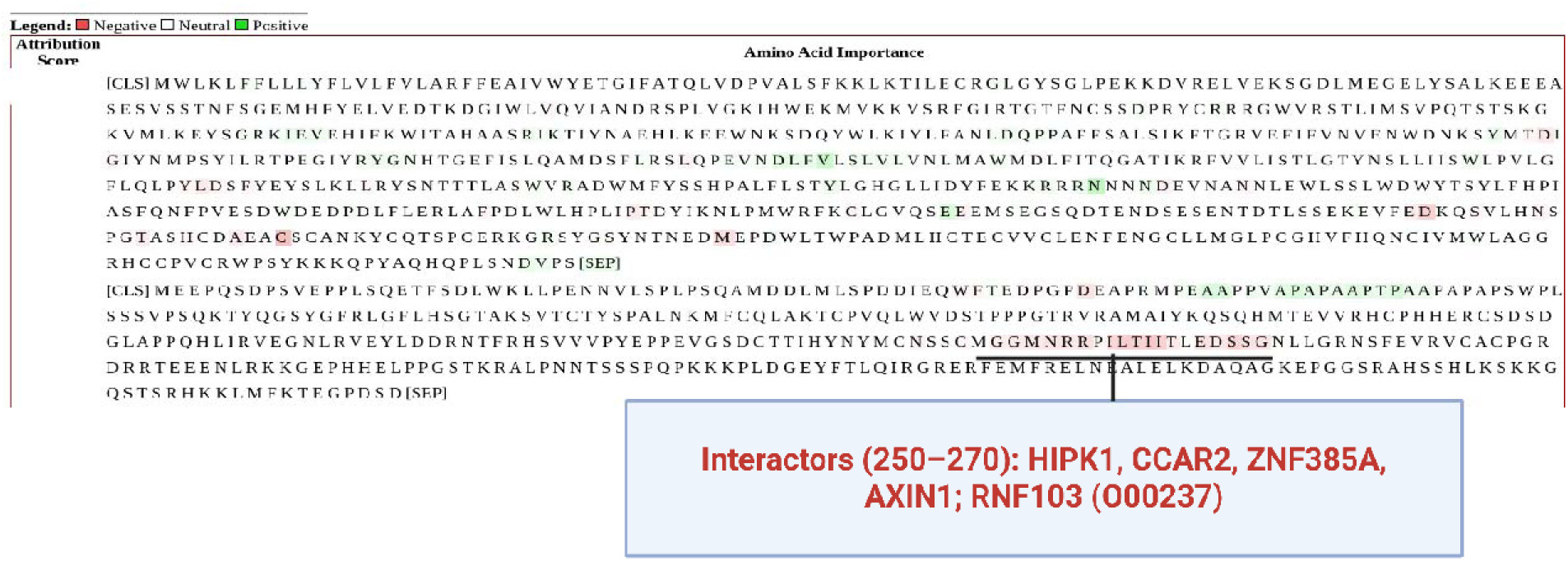
**- Interactors (250–270): HIPK1, CCAR2, ZNF385A, AXIN1; RNF103 (O00237) linked to p53 R175H**

The model assigns a positive attribution to the trans-membrane segment of RNF103 (residues 320–330, Figure 4 <u>a</u>), which serves as an ER anchor that correctly orients the ligase for ubiquitinating cytosolic substrates such as p53 [85]. In DTX2, substrate recognition relies on both flexible and ordered modules: the positively scored intrinsically disordered linker (residues 350–390, Figure 5 <u>a</u>) can adapt to the proline-rich 60–85 (Figure 4 <u>c</u>) stretch of p53 via short linear motifs [86], whereas the C2H2-type zinc-finger (residues 412–473, Figure 5 <u>b</u>) offers a well-folded surface that complements p53 features and reinforces binding specificity [87]. Together these elements illustrate how IDRs supply adaptability while structured domains provide stabilizing contacts [84].

**Figure 5.**
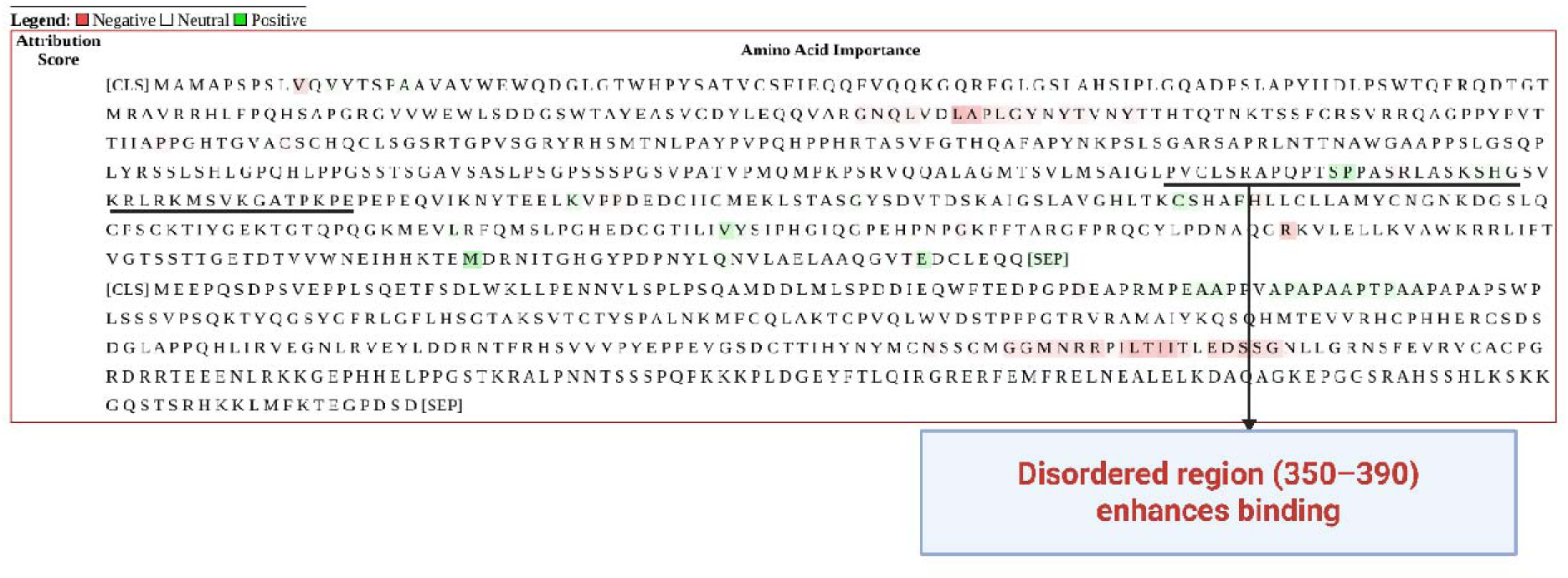
a - **Disordered region (350–390) enhances binding; Interpretation of Q86UW9 (DTX2) and p53 R175H**

**Figure 5b.**
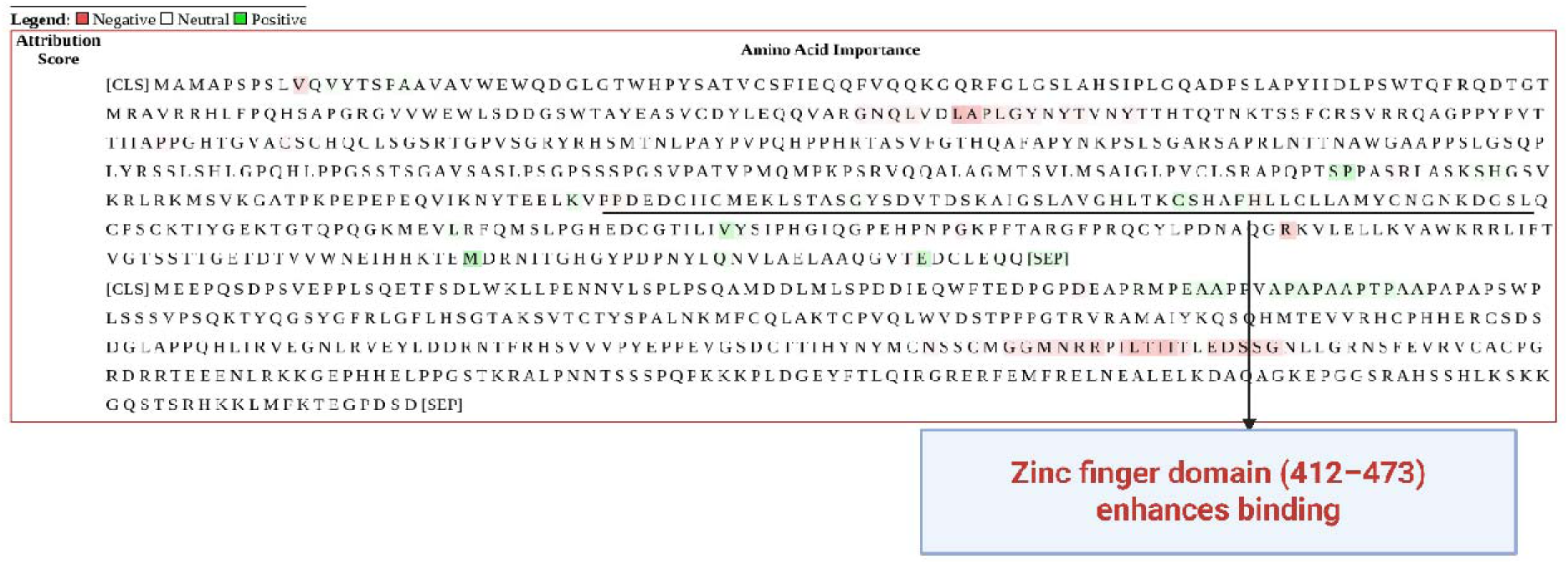
**- Zinc finger domain (412–473) enhances binding, Interpretation of Q86UW9 (DTX2) and p53 R175H**

By contrast, the negative attribution assigned to the acidic, disordered loop of RNF103 (residues 550–570, Figure 4 <u>b</u>) likely reflects an autoinhibitory mechanism: in many RING-type E3 ligases, an internal acidic segment can fold back over the RING domain, shielding catalytic cysteines or the E2-binding groove until a regulatory signal-often phosphorylation-displaces it. In such a closed conformation the loop would dampen, rather than promote, p53 ubiquitination-explaining its negative weight in the attribution map.

Integrated gradients indicate that RNF103 and DTX2 favor disordered, flexible regions of p53 that enable adaptable engagement. While ordered, structured segments are penalized when their geometry/chemistry mismatches the ligase surface. The ligases appear to use two mode strategy-flexible elements to initiate capture and folded modules to stabilize specificity. Negative attributions map to autoinhibitory features, consistent with regulated activation of E3 activity rather than constitutive binding.

### 3.6. Example application 2: Eliminating an undruggable pathogenic structural protein in neurodegenerative diseases

Structural proteins are a small yet crucial class of potential drug targets (∼2-3% of approved drugs). Their poor druggability derives from a lack of deep pockets, ubiquitous expression, and the risk of severe side-effects when perturbed. Paclitaxel illustrates a successful exception, binding tubulin to stabilize microtubules and arrest mitosis in cancer therapy [88,89,90]. Another structural protein, Tau (encoded by MAPT), is essential for microtubule stability in neurons. The P301L Tau variant disrupts the microtubule-binding domain, exposes hydrophobic regions that nucleate neurofibrillary tangles, and underlies frontotemporal dementia (FTD) . The corresponding UBE4B and UBR1 attribution maps are presented in Figure 6a, Figure 6b, Figure 6c, Figure 6d, Figure 6e, Figure 6f, Figure 7a, Figure 7b, and Figure 7c.

**Figure 6.**
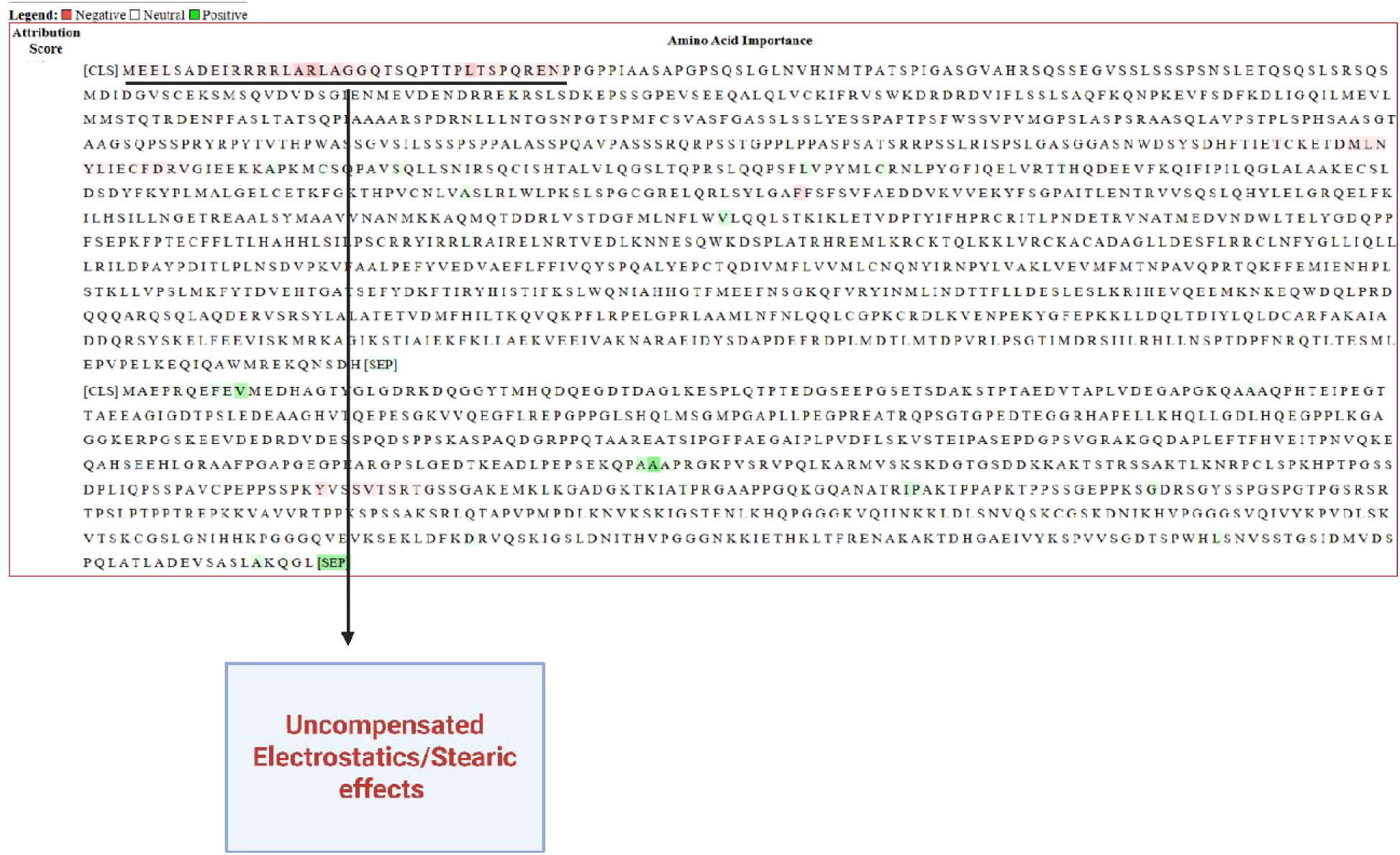
a - **Electrostatic/steric effects (1–40), Interpretation of O95155 (UBE4B) and Tau**

**Figure 6b.**
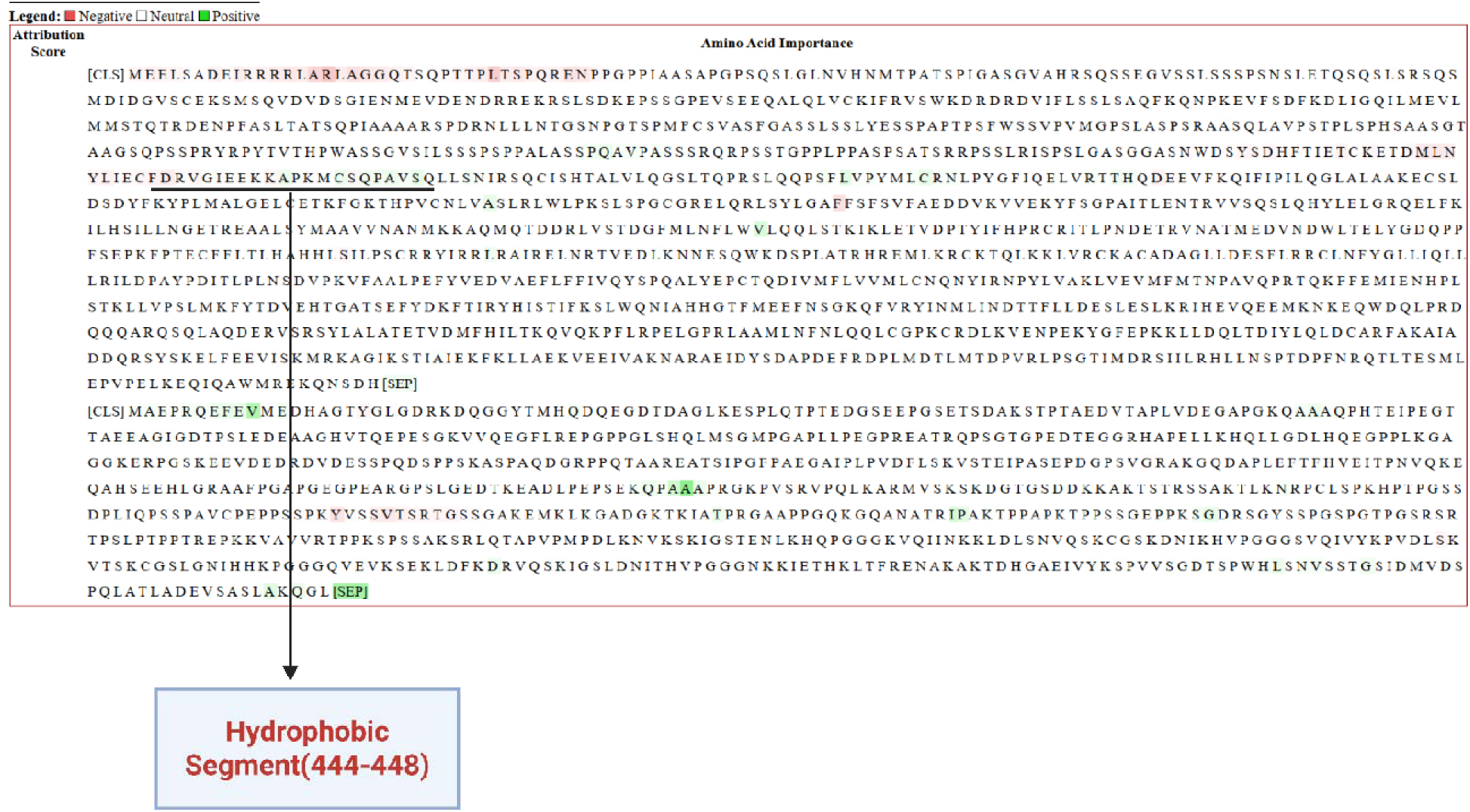
**- Hydrophobic segment, Interpretation of O95155 (UBE4B) and Tau**

**Figure 6c.**
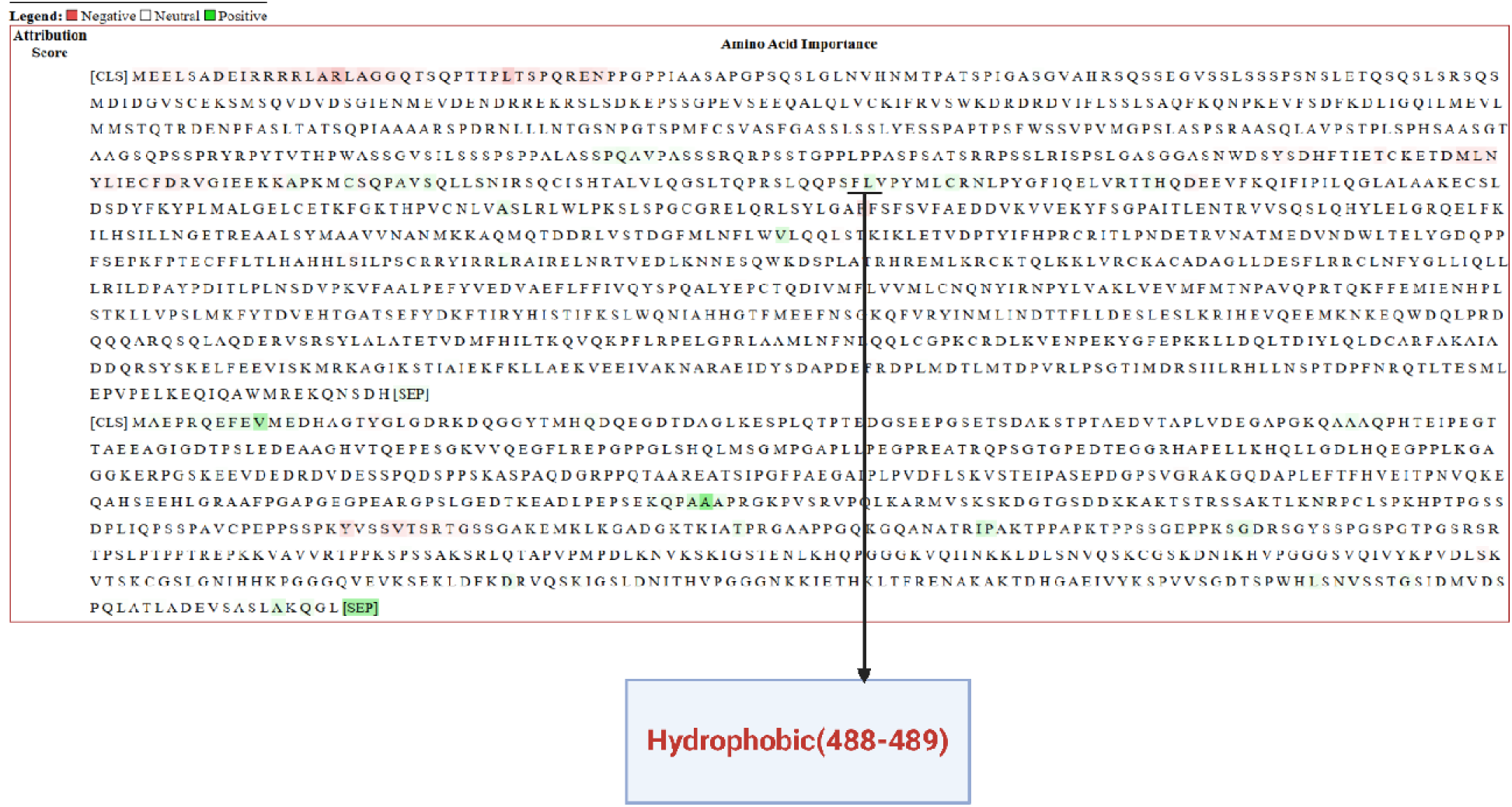
**– Another Hydrophobic segment, Interpretation of O95155 (UBE4B) and Tau**

**Figure 6d.**
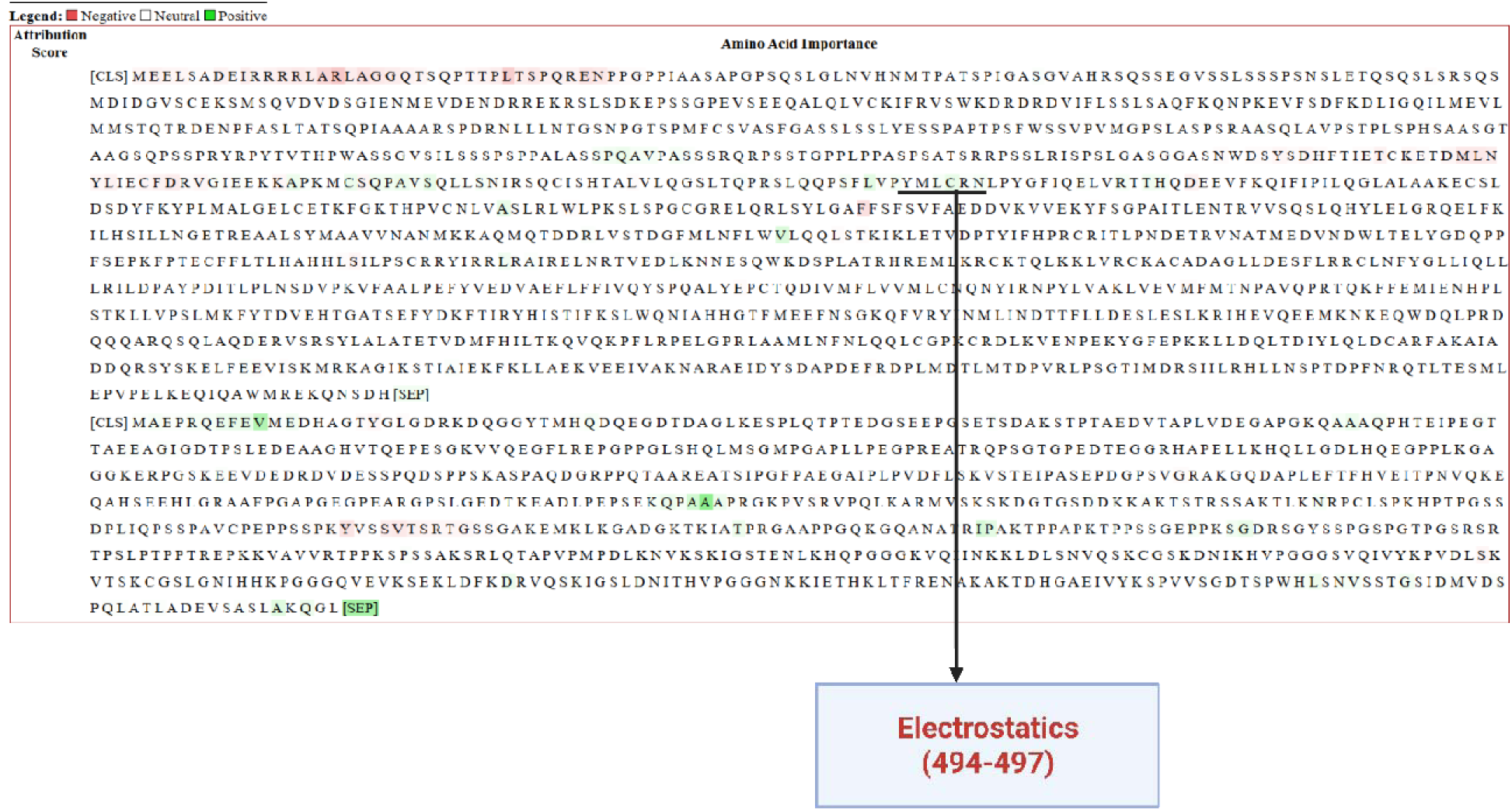
**– Electrostatic Interactions, Interpretation of O95155 (UBE4B) and Tau**

**Figure 6e.**
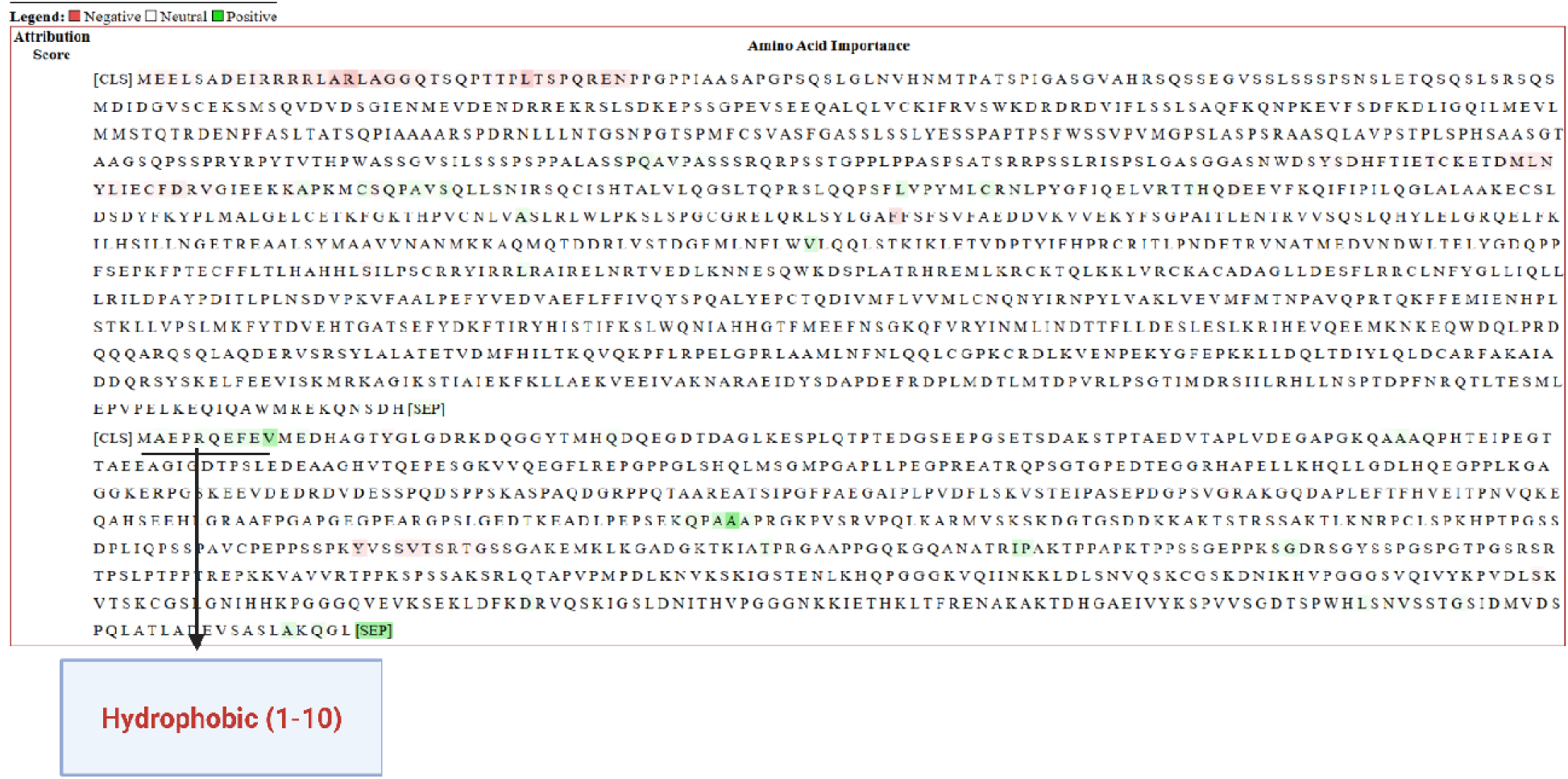
**– Hydrophobic Residues in Tau, Interpretation of O95155 (UBE4B) and Tau**

**Figure 6f.**
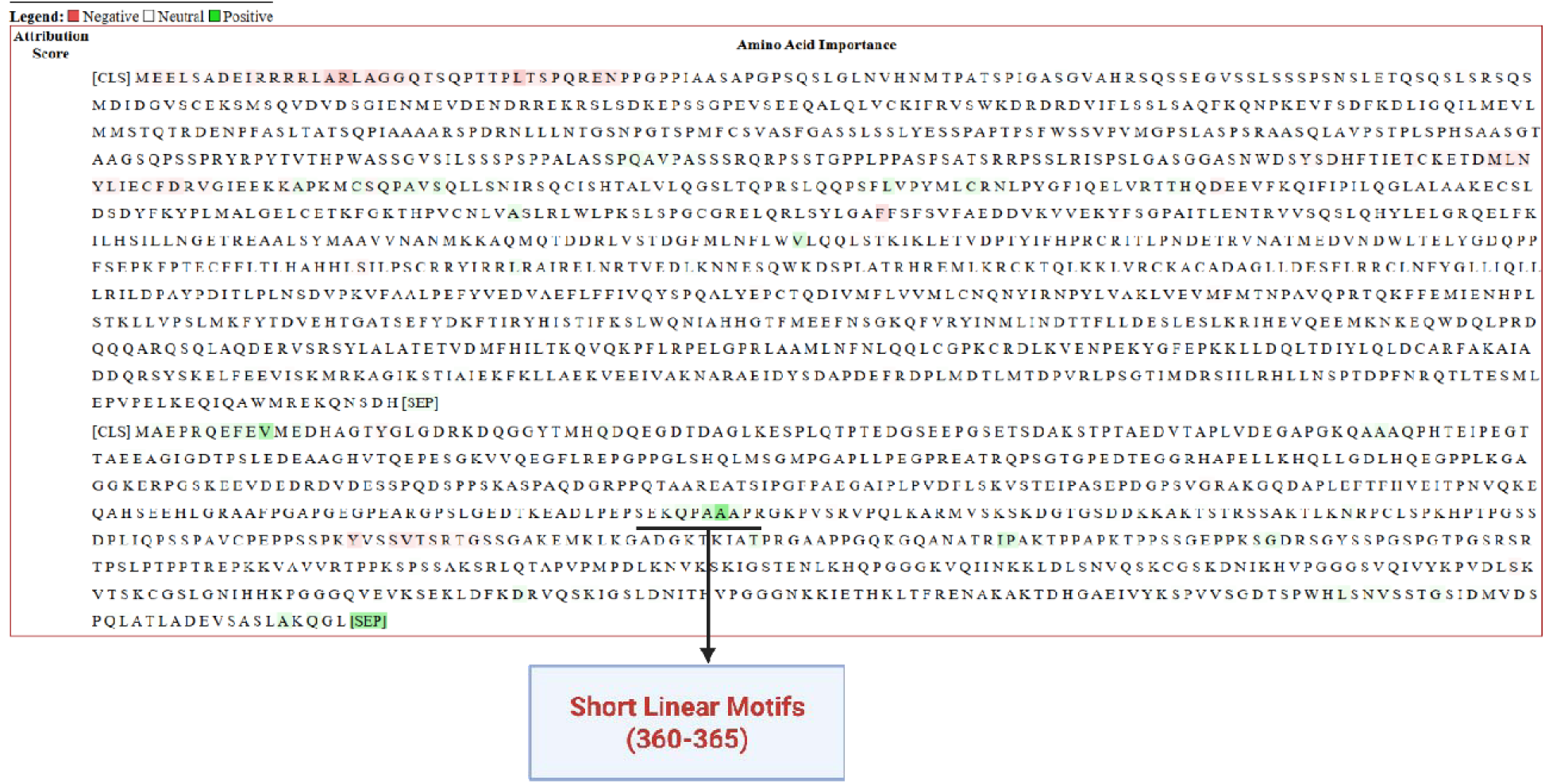
**– Short Linear Motifs in Tau, Interpretation of O95155 (UBE4B) and Tau**

**Figure 7.**
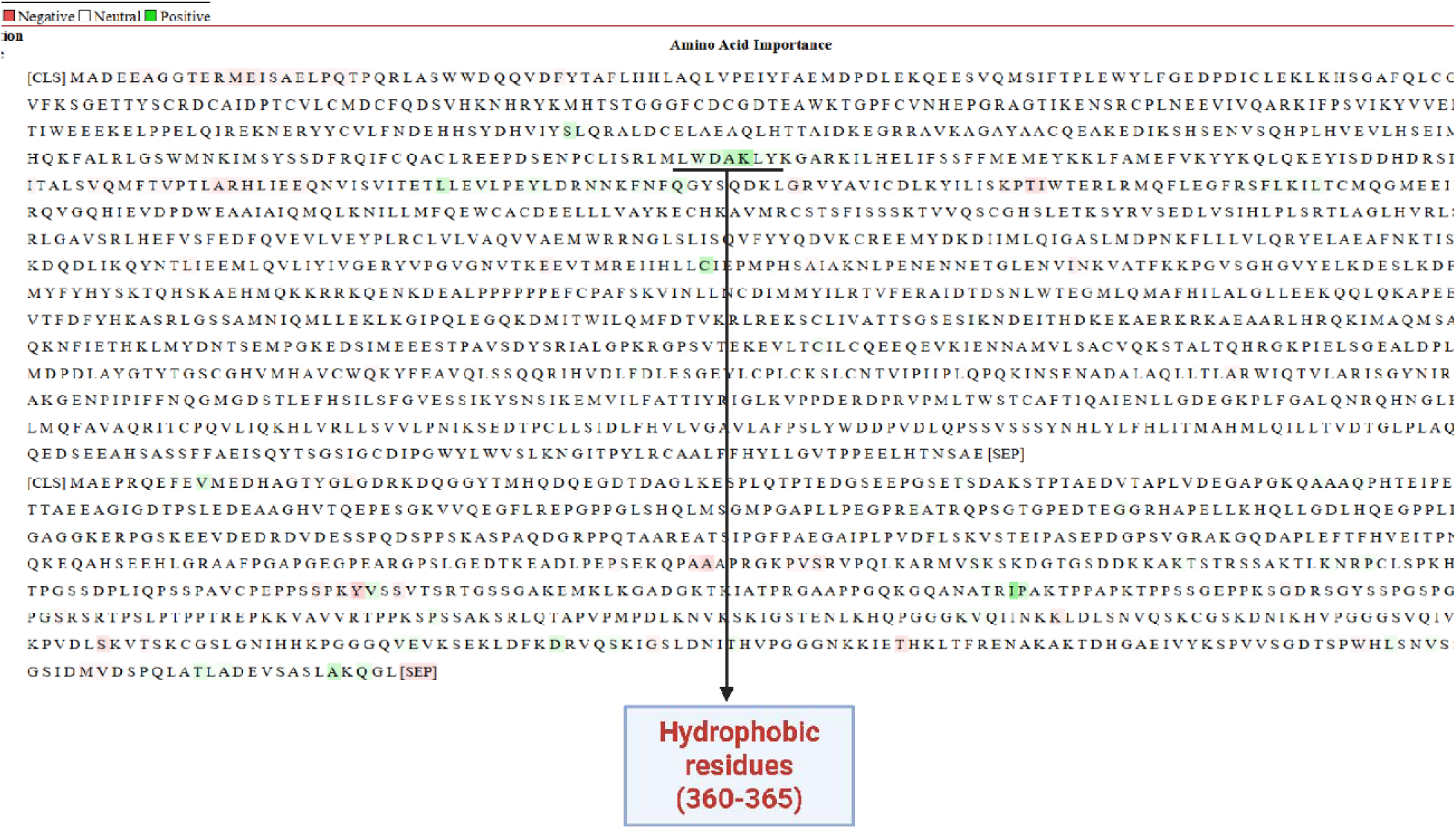
a – **Hydrophobic residues, Interpretation of Q8IWV7 (UBR1) and Tau**

**Figure 7b.**
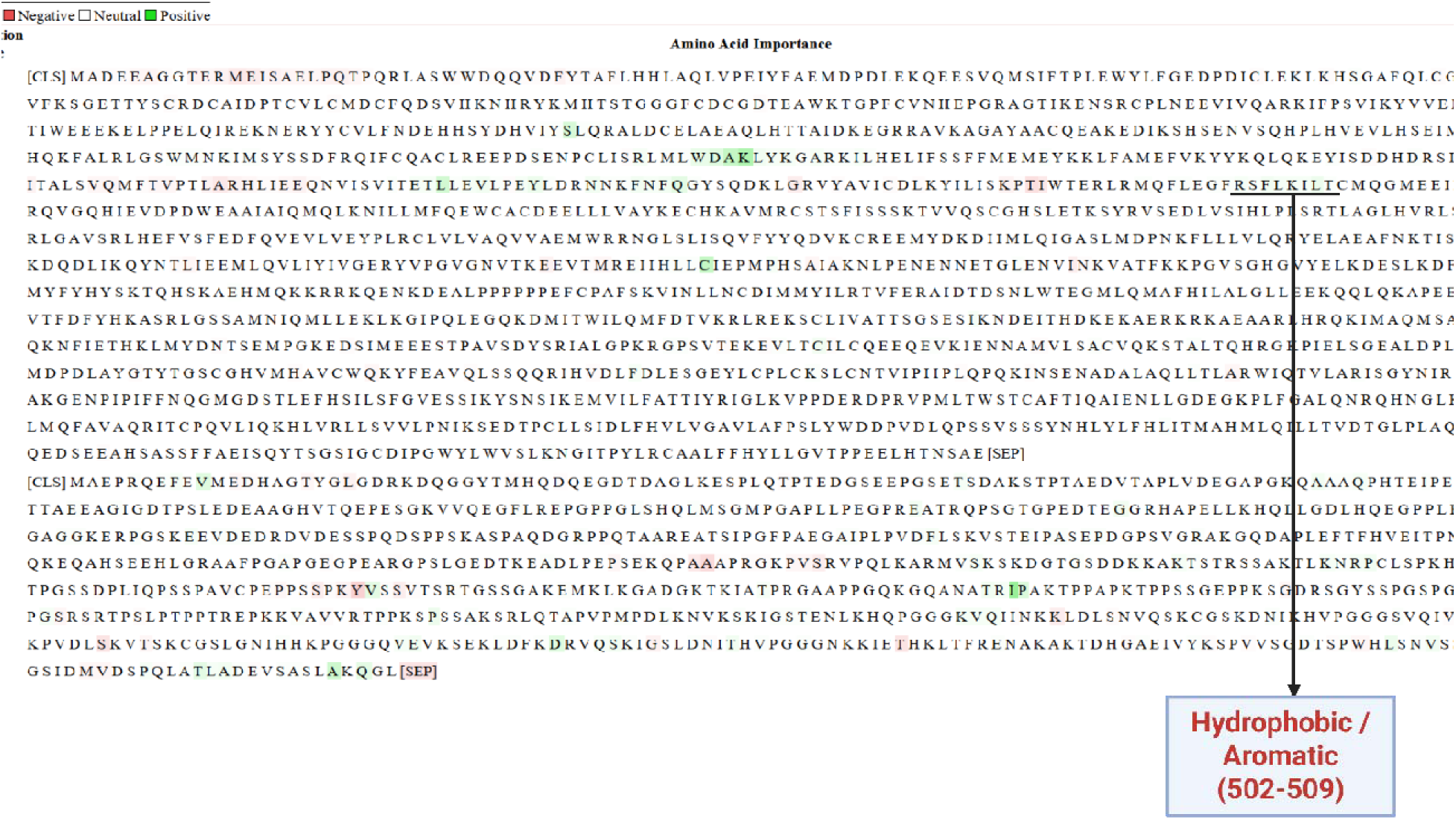
**– Hydrophobic and Aromatic Residues, Interpretation of Q8IWV7 (UBR1) and Tau**

**Figure 7c.**
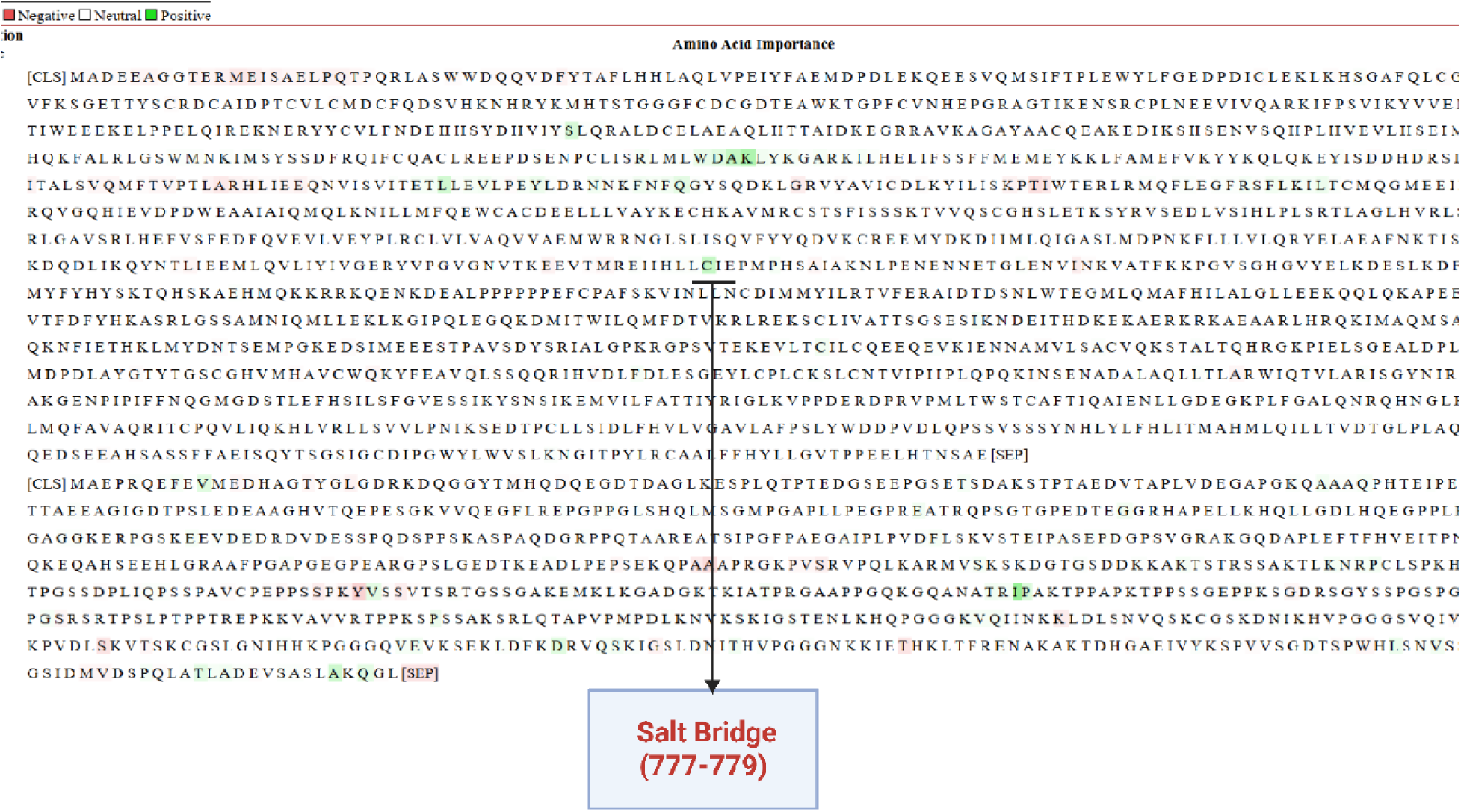
– Salt Bridge residues on Ligase, Interpretation of Q8IWV7 (UBR1) and Tau

Neurodegenerative diseases represent a major and growing public health concern, with rising incidence rates—particularly in Western countries. Among these disorders, Alzheimer’s disease and frontotemporal dementia (FTD) together account for approximately 80% of cases, with the microtubule-associated protein Tau playing a central pathological role in both [91]. A well-characterized pathogenic variant, Tau P301L, has been strongly implicated in the development of both Alzheimer’s disease and FTD, where it promotes neurodegeneration through enhanced aggregation, reduced microtubule binding, and increased cytotoxicity [92,93,94,95,96,97]. In this study, our model predicts a ranked list of E3 ligases that may participate in tissue-specific interactions involving mutant Tau as shown in Table 6 , this information can be used for drug discovery.

**Table 6.**
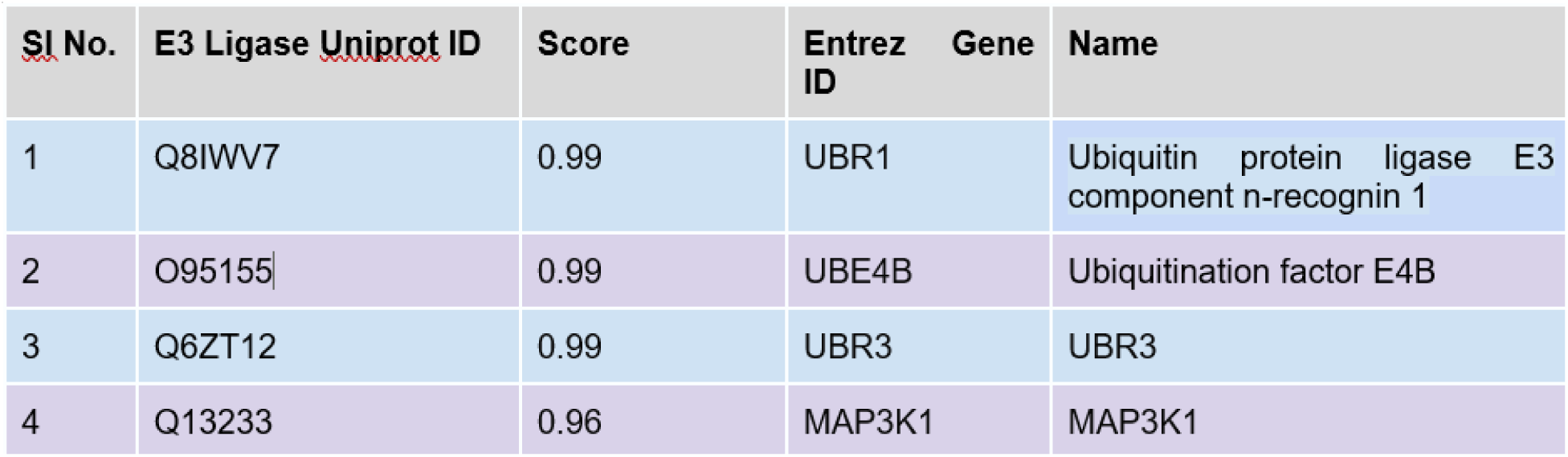
- Predictions - Ranked E3Ls in Tau mutation in Alzheimer’s disease.

It is important to note that while the predicted protein–protein interactions are tissue-specific, this does not necessarily imply that both the E3 ligase and the substrate (e.g., Tau P301L) are tissue–specific. It is important to acknowledge that many of the E3 ligases are not inherently tissue specific [16,31]. Therefore, we conduct an independent tissue-specificity analysis of the E3 ligases, drawing from multiple data sources to determine the target tissue of the predicted ligase, as discussed in subsequent sections [98,99].

In line with the p53 use case, we employed Integrated Gradients (IG) to explain the model’s predicted interactions of Tau with UBR1 and UBE4B. Figures 6a, <u>6b</u>, <u>6c</u>, <u>6d</u>, <u>6e</u>, <u>6f</u>, <u>7a</u>, <u>7b</u>, and <u>7c</u> map residue-level attribution scores for the interactions between the E3 ligases UBE4B and UBR1 and the Alzheimer’s-associated Tau P301L variant, with red and green intensities marking residues that respectively promote or hinder binding The IG analysis indicates that the model’s predictions are grounded in sound biophysical principles. Specifically, the hydrophobic and polar amino acids located at positions 1-10 (Figure 6 <u>e</u>; [100]) in Tau form a canonical hotspot interaction motif that contributes significantly to the free energy of binding. This is followed by short linear motifs at positions 360-365 (Figure 6 <u>f</u>; [101]) which play a key role in interactions with both ligases under discussion.

For UBR1, hydrophobic and aromatic residues at positions 360-365 (Figure 7 <u>a</u>; [100]) and 502– 509 (Figure 7 <u>b</u>) are predicted to contribute to Tau binding through tight van der Waals packing and electrostatic interactions involving R/K residues, collectively enhancing complex stability. Finally, polar residues at positions 777-779 (Figure 7 <u>c</u>) are implicated in forming electrostatic and hydrogen bonds with complementary Tau residues, providing additional stabilization to the UBR1–Tau complex [102].

A similar hydrophobic and polar amino acid pattern is observed in UBE4B at positions 444-448 (Figure 6 <u>b</u>), with additional hydrophobic contacts at positions 488-489 (Figure 6 <u>c</u>). Polar and charged groups at positions 494-497 (Figure 6 <u>d</u>) further suggest hydrophobic packing and potential salt bridge formation in UBE4B–Tau interactions. The negative attribution observed in UBE4B across positions 1-40 (Figure 6 <u>a</u>) aligns with the presence of uncompensated electrostatic interactions and steric hindrance [103].

Table 4, Table 5, Table 6 and Table 7 summarize the predicted E3-ligase candidates-reporting their interaction scores (0-1), sub-cellular localizations, and genomic annotations for gastrointestinal cancer and Alzheimer’s disease, respectively. These predictions are potential hypotheses for further validation.

**Table 7.**
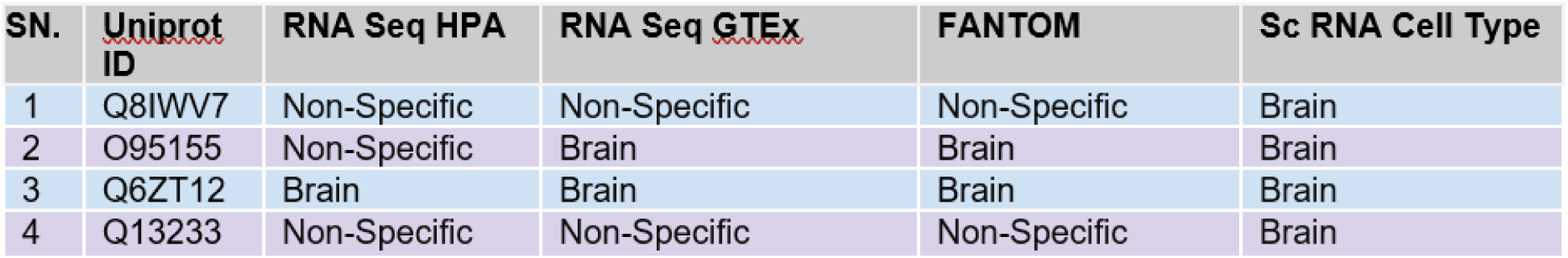
- Tissue Categorization based on Bulk/Single RNA Seq of E3 Ligases

## 4. Limitations and future directions

ELITE is built upon the strong theoretical foundations of attention-based models, which are well suited for capturing long-range dependencies in biological sequences. However, its performance is constrained by the limited availability of data for task-specific fine-tuning. Although we employ a sophisticated architecture and leverage tissue-resolved datasets for training, the average size of tissue-specific datasets typically lies in the hundreds of samples, which can restrict achievable performance at inference time. Integrating diverse and heterogeneous data sources at scale represents a promising strategy to alleviate data scarcit and further enhance model robustness and predictive accuracy.

the landscape of PLMs, the foundation layer of ELTE is rapidly evolving terms of data and training paradigms. The E3 Ligase prediction problem stand to benefit directly from these scientific and technological advancements. Beyond ProtBERT newer models tend to capture protein–protein interactions (PPIs) better because they either (i) learn richer long-range context from larger-scale pretraining, (ii) encode stronger structure/interface-relevant signals, or (iii) add reasoning modules explicitly designed for interaction prediction. Within the ProtTrans family, ProtT5 (T5-style PLM) is often a stronger embedding backbone than ProtBERT, with multiple studies reporting improved downstream behavior when swapping ProtBERT embeddings for ProtT5 embeddings (including interaction/binding-related predictors) [35]. In parallel, ESM-2 improves on earlier encoder LMs by scaling and training refinements that yield embeddings with stronger emergent structural information—useful for PPIs because interfaces are fundamentally structural objects [104]. MSA Transformer goes further by taking multiple sequence alignments as input and leveraging explicit evolutionary context, which can sharpen interface/contact signals relative to single-sequence models like ProtBERT (at the cost of requiring MSAs) [42]. Finally, "new" PPI-specific architectures such as PPITrans are typically better than a ProtBERT classifier baseline because they don’t just embed proteins independently—they learn interaction-aware cross-protein representations, improving generalization (e.g., across species) in reported evaluations [105].

Beyond Siamese sequence encoders, structure-aware geometric and hybrid PLM–graph architectures typically provide a stronger inductive bias for PPI prediction because they explicitly model interactions in the 3D space where binding occurs [106,107]. Geometric deep learning approaches represent each protein as a spatial object (e.g., surface/point cloud/graph) and use equivariant or geometry-attentive message passing to learn interface features that depend on shape complementarity, local chemistry, and spatial proximity—signals that are difficult for Siamese embeddings to recover from sequence alone [106,108,109]. Interface-focused geometric models such as PInet and geometric-transformer contact predictors (e.g., DeepInteract’s Geometric Transformer) are concrete examples of this structure-first trend [106]. Building on this, hybrid frameworks fuse protein language model (PLM) residue embeddings with structural graphs (nodes = residues/atoms; edges = spatial neighbors), so that the PLM contributes evolutionary/functional context while the graph/geometry module enforces physically meaningful locality [107,110]. This combination is repeatedly reported to improve robustness across protein–protein tasks (including PPI/interface/contact prediction), and it is increasingly emphasized in recent inter-protein contact and structure-aware PLM work as a high-performing direction [111]

## 5. Conclusion

ELITE harnesses the representational power of large protein language models (PLMs) to capture intricate sequence patterns and transfer them effectively to downstream tasks. The dense, semantically rich embeddings generated by these PLMs not only separate interacting from non-interacting pairs with high fidelity but also prioritize sensitive detection of the positive class-i.e., true protein–protein interactions.

The shared-weight Siamese network encodes each protein independently within its tissue context and combines their representations into a single product tensor, enabling the capture of highly non-linear sequence dependencies that govern complex and poorly characterized protein–protein interactions. Integrated Gradients applied to high-confidence predictions reveal key residue-level features contributing to interaction likelihood, with results supported by existing literature. With a strong theoretical foundation, interpretable outputs, and a modular design, this architecture offers a powerful and adaptable framework that can be efficiently fine-tuned to tissue-specific or custom PPI datasets.

As a demonstrative use case, we deployed our model to identify tissue-specific E3 ubiquitin ligases suitable for PROTAC design against pathogenic variants of two classically intractable proteins, p53 and Tau, which drive gastrointestinal cancers and neurodegenerative disorders such as Alzheimer’s disease and frontotemporal dementia. The model produced high-confidence ligase rankings for each disease-relevant tissue, and Integrated Gradients attribution traced the sequence features underpinning every prediction, affording mechanistic transparency. These ligase–substrate hypotheses constitute strong leads for experimental validation and lay the groundwork for variant-targeted PROTAC therapeutics.

Beyond the scope of the study, it would also be interesting to explore variant selectivity by evaluating whether mutations in the target expose or alter motifs that render the mutant—but not the wild-type—amenable to ligase engagement and degradation.

To our knowledge, this is the first study to leverage large language models for E3 ligase selection in targeted protein degradation, offering an innovative solution to longstanding hurdles of sparse data, biochemical complexity, and limited model interpretability. The architecture’s modular design and flexible classifier allow it to accommodate datasets of widely varying sizes and noise, enabling straightforward adaptation to new targets and experimental contexts.

## Resource availability

### Lead contact

Further information and requests for code and data should be directed to and will be fulfilled by the lead contact, Holger Fröhlich.

## Materials availability

This study did not generate new unique reagents.

## Data and code availability

Code and data are located at: https://github.com/sabyasachi2k13/ELITE

Any additional information required to reanalyze the data reported in this paper is available from the lead contact upon request.

## Declaration of competing interest

The authors declare no competing interests related to this study.

## CRediT authorship contribution statement

Conceptualization, H.F and S.P.; Methodology, H.F. and S.P.; Data Curation, Formal Analysis, Visualization, Investigation, Validation, S.P.; Technical Help: S.M; Supervision, H.F; Writing— Original Draft, S.P.; Writing—Review and Editing, S.P and H.F.

## Acknowledgments

We thank André Gemünd for his support regarding the computational infrastructure of Fraunhofer SCAI.

## Data science maturity

DSML3: Development/pre-production: Data science output has been rolled out/validated across multiple domains/problems

## Supplementary material

Document S1

## Declaration of generative AI and AI-assisted technologies in the writing process

The authors utilized OpenAI’s GPT-3o only to enhance the clarity of the text and thoroughly reviewed the content to ensure its scientific accuracy.

## Notes

### Competing Interest Statement

The authors have declared no competing interest.

### Summary of Updates

added new models, ESMC ESM 650 M in addition to Protbert

