## Supplementary for "ELITE: E3 Ligase Inference for Tissue specific Elimination: A LLM Based E3 Ligase Prediction System for Precise Targeted Protein Degradation"

Supplementary Material - Document S1

Tissue aggregation, balanced evaluation, tissue-specific E3-ligase assignment, and training details.

S1. Data mapping

Table S1. Functional aggregation of tissues into 26 tissue groups.

| **Sl** | **Tissue** | **Tissue** | **Tissue** | **Tissue** | **Tissue** | **Tissue** | **Group** |
| --- | --- | --- | --- | --- | --- | --- | --- |
| **1** | Adrenal Cortex | Adrenal Gland |  |  |  |  | **Adrenal** |
| 2 | Cochlea | Ear |  |  |  |  | **Auditory** |
| 3 | Blood | Blood Plasma | Blood Platelet | Serum |  |  | **Blood** |
| 4 | Bone | Bone Marrow | Osteoblast |  |  |  | **Bone** |
| 5 | Amygdala | Astrocyte | Basal Ganglion | Brain | Caudate Nucleus | Hypothalamus | **Brain** |
|  | Caudate Putamen | Central Nervous System | Cerebellar Cortex | Cerebellum | Cerebral Cortex | Locus Coeruleus |  |
|  | Corpus Callosum | Corpus striatum | Dentate Gyrus | Diencephalon | Forebrain | Medulla Oblongata |  |
|  | Forebrain | Frontal Lobe | Glia | Hippocampus | Midbrain | Nervous System |  |
|  | Nucleus Accumbens | Occipital Lobe | Occipital Pole | Parietal Lobe | Parietal Lobe | Pons |  |
|  | Substantia Nigra | Subthalamic Nucleus | Telencephalon | Temporal Lobe | Thalamus |  |  |
| 6 | Aorta | Artery | Cardiac Muscle | Heart |  |  | **Cardiac** |
| 7 | Adipose | Muscle | Smooth Muscle | Cardiac Muscle |  |  | **Muscle** |
| 8 | Adrenal Cortex | Adrenal Gland | Hypophysis |  |  |  | **Endocrine** |
| 9 | Embryo | Fetus | Placenta | Trophoblast | Umbilical Cord | Umbilical Vein Endothelial Cell | **Fetal** |
| 10 | Cecum | Colon | Duodenum | Esophagus | GI Tract | Esophagus | **GastroIntestinal** |
|  | GastroIntestinal Tract | Ileum | Intesting | Jejunum | Large Intestine | Liver |  |
|  | Pancreas | Pancreatic Islet | Small Intestine | Stomach | Vermiform Appendix |  |  |
| 11 | Hepatocyte |  |  |  |  |  | **Hepatic** |
| 12 | B Lymphocyte | Basophil | CD8 Cell | Dendritic | Eosinophil | Granulocyte | **Immune** |
|  | Hematopoietic Stem cell | Leukocyte | Lymph Node | Lymphocyte | Macrophage | Mast Cell |  |
|  | Megakaryocyte | Monocyte | Mononuclear  Phagocyte | NK Cell | Spleen | T Lymphocyte |  |
|  | Spleen | Lymphocyte | Thymocyte | Tonsil |  |  |  |
| 13 | Mammary Epithelium | Mammary Gland |  |  |  |  | **Mammary** |
| 14 | Salivary Gland | Tooth |  |  |  |  | **Oral** |
| 15 | Peripheral Nervous System |  |  |  |  |  | **PNS** |
| 16 | Bronchial Epithelial Cell | Bronchus | Lung | Trachea |  |  | **Pulmonary** |
| 17 | Kidney | Nephron | Podocyte | Renal Glomerulus | Renal Tubuie |  | **Renal** |
| 18 | Corpus Luteum | Myometrium | Ovarian Follicle | Ovary | Oviduct | Uterine Cervix | **Reproductive**  **Female** |
|  | Uterine Endometrium | Uterus |  |  |  |  |  |
| 19 | Prostate Gland | Spermatid | Spermatocyte | Spermatogonium | Testis |  | **Reproductive**  **Male** |
| 20 | Cartilage | Chondrocyte | Skeletal Muscle |  |  |  | **Bone and Skeletal Muscle** |
| 21 | Epidermis | Hair Follicle | Keratinocyte | Skin | Skin Fibroblast |  | **Skin** |
| 22 | Nervous System | Spinal Cord |  |  |  |  | **Spinal Cord** |
| 23 | Thyroid Gland |  |  |  |  |  | **Thyroid** |
| 24 | Urinary Bladder | Uroepithelium |  |  |  |  | **Urinary Tract** |
| 25 | Blood Vessel | Vascular Endothelial Cell | Vascular Endothelium |  |  |  | **Vascular** |
| 26 | Choroid | Cornea | Eye | Lens | Retina | Tear gland | **Vision** |

S2. Testing under balanced positive-to-negative ratio

Table S2. Performance under a balanced positive-to-negative ratio. All values are percentages.

| **Sl.** | **Tissue** | **PR-AUC** | **Precision** | **Recall** | **Accuracy** |
| --- | --- | --- | --- | --- | --- |
| 1 | Cardiac | 99 | 90 | 99 | 90 |
| 2 | Adrenal | 99 | 99 | 98 | 98 |
| 3 | Bone | 99 | 99 | 99 | 96 |
| 4 | Brain | 99 | 95 | 97 | 93 |
| 5 | Renal | 88 | 89 | 99 | 89 |
| 6 | PNS | 94 | 90 | 99 | 90 |
| 7 | Immune | 99 | 96 | 97 | 94 |
| 8 | Auditory | 94 | 92 | 99 | 92 |
| 9 | Blood | 99 | 93 | 99 | 93 |
| 10 | Reproductive F | 99 | 97 | 98 | 96 |
| 11 | Mammary | 99 | 99 | 97 | 96 |
| 12 | Connective | 99 | 93 | 99 | 93 |
| 13 | Endocrine | 99 | 95 | 99 | 95 |
| 14 | Fetal | 99 | 96 | 99 | 95 |
| 15 | Gastrointestinal | 98 | 91 | 99 | 92 |
| 16 | Hepatic | 95 | 90 | 99 | 90 |
| 17 | Oral | 99 | 98 | 97 | 96 |
| 18 | Pulmonary | 87 | 89 | 99 | 98 |
| 19 | Reproductive M | 99 | 95 | 99 | 95 |
| 20 | Skeletal | 97 | 94 | 99 | 94 |
| 21 | Skin | 99 | 98 | 97 | 96 |
| 22 | Spinal Cord | 99 | 95 | 99 | 95 |
| 23 | Thyroid | 99 | 95 | 99 | 95 |
| 24 | Urinary Tract | 99 | 97 | 99 | 97 |
| 25 | Vascular | 99 | 99 | 99 | 95 |
| 26 | Vision | 99 | 95 | 99 | 94 |

S3. Tissue-specific assignment of E3 ligases

To prioritize E3-ligase candidates for experimental validation, we evaluated each predicted ligase systematically. Because all candidates display high in-silico binding affinity to the target substrate, tissue and cell-type specificity became the critical discriminant [1]. Determining true enrichment is non-trivial: expression must be inferred indirectly from multiple modalities—bulk and single-cell RNA-seq, protein levels by immunohistochemistry, and cDNA abundance—each curated in the Human Protein Atlas (HPA) [1-4]. We validated the transcriptomic and proteomic signatures corresponding to each predicted E3 ligase by integrating data from multiple sources. This comprehensive analysis enabled us to assign a definitive tissue or cell-type context to each ligase, thereby reinforcing the biological relevance of our predictions and refining the E3 Ligase list for downstream validation.
